# Exercise-Induced Myostimulin Enhances Muscle Function in Health and Disease

**DOI:** 10.1101/2025.04.09.647960

**Authors:** Hillger Frank, Keller Hansjörg, Furrer Angela, Melly Stefan, Bösch Julian, Seddik Sabrina, Pierrel Eliane, Bello Fiona, Scharenberg Meike, Leber Xavier, Hug Christian, Vicart Axel, Greiner Geraldine, Meyer Angelika, Cobos-Correa Amanda, Kaufmann Markus, Aschenbrenner Dominik, Allard Cyril, Delucis-Bronn Corinne, Killmer Saskia, Hils Lukas, Rothen Julian, Lemire Sophie, Altorfer Marc, Lambert Christian, Doelemeyer Arno, Accart Nathalie, Bepperling Alexander, Weiss Laurent, Burget Aurelia, Couttet Philippe, Parker N. Christian, Rooks Daniel, Kneissel Michaela, Summermatter Serge

**Affiliations:** Biologics Research Center, Novartis Biomedical Research, Novartis Pharma AG, Basel, Switzerland; Diseases of Aging and Regenerative Medicine, Biomedical Research, Novartis Pharma AG, Basel, Switzerland; Preclinical Safety, Novartis Biomedical Research, Novartis Pharma AG, Basel, Switzerland; Discovery Sciences, Novartis Biomedical Research, Novartis Pharma AG, Basel, Switzerland; Immunology Disease Area, Biomedical Research, Novartis Pharma AG, Basel, Switzerland; Sandoz, Biodevelopment, Holzkirchen, Germany; Translational Medicine, Biomedical Research, Novartis Pharma AG, Cambridge, United States

**Keywords:** exercise mimetic, myokine, skeletal muscle, muscle strength, satellite cells, myostimulin

## Abstract

Musculoskeletal diseases are a leading contributor to years lived with disability worldwide^1,2^. While exercise offers significant benefits for people with these conditions, many individuals do not engage in adequate physical activity^3^. Consequently, there is growing interest in pharmacological interventions that can emulate essential health-promoting effects of exercise^4,5^. By integrating transcriptomics data of exercised skeletal muscle, we identified *C1orf54/*C1ORF54 as a novel exercise-responsive gene in mice and humans. We demonstrate that removal of the first sixteen N-terminal amino acids of C1ORF54 gives rise to a previously uncharacterized protein that stimulates the proliferation of muscle precursor cells and which we named myostimulin. Intriguingly, repeated intermittent treatment of mice with recombinant myostimulin boosts maximal isometric strength in mice within a week. Moreover, we have engineered a variant with improved biophysical properties, increased biological activity *in vitro* and enhanced efficacy *in vivo*. This variant even accelerates the recovery of muscle strength from axonotmesis, a condition associated with pronounced muscle weakness. Our data ascribe to myostimulin a role for enhancing the regenerative capacity of skeletal muscle and mediating functional adaptations characteristic of sustained resistance training. Therefore, myostimulin could be an innovative, fast acting therapeutic for certain human musculoskeletal diseases, injuries and other disorders that improve with exercise.

## MAIN

Many common acute and chronic diseases are associated with musculoskeletal ailments. For instance, patients with disuse atrophy post-surgery, lower extremity injuries, neurological disorders, or age-related health restrictions often experience a progressive loss of muscle mass and function^6–8^. These impairments in muscle integrity have wide adverse consequences, such as severe functional limitations, an increased risk of falling, and reduced quality of life^6–8^. Musculoskeletal conditions are prevalent in men and women of all ages across all socio-demographic strata of our modern society and affect hundreds of millions of people globally^6,8^. The prevalence of these conditions rises with age. Accordingly, population ageing, driven by increasing longevity, and the progression of large-sized cohorts to older ages, lead to a steadily growing number of people with musculoskeletal impairments. This problem is further accentuated by the sedentary lifestyle prevailing in our modern society. Therefore, these conditions represent increasing unmet medical and societal needs.

Exercise is a well-recognized and highly efficacious therapy for musculoskeletal diseases^9,10^. In fact, there is evidence for exercise as therapy for at least 26 different chronic diseases involving musculoskeletal impairments and other disorders that can indirectly aggravate musculoskeletal conditions^9^. However, it is only partially established how exercise mediates its health-promoting effects^11^. Some of the beneficial adaptations to exercise seem to be governed by muscle-derived proteins, which are secreted by contracting skeletal muscles and are referred to as myokines^12,13^. Such myokines exert their actions in an endo-, auto-or paracrine fashion and can provide a feedback loop for muscle tissue to self-regulate its adaptation to exercise training^12–14^. A prominent adaptation regulated by myokines is the activation of muscle-resident stem cells, which play a key role in the growth, remodeling and regeneration of skeletal muscle tissue.

Problematically, many people are unable to reach the levels of physical activity required to maintain muscle health; in particular patients who cannot exert strenuous physical activities because of musculoskeletal impairments^3^. While there are anabolic agents that increase muscle mass, such substances can result in a variety of undesired side effects. For instance, anabolic steroid use has been associated with high blood pressure^15^, decreased function of the heart’s ventricles^16^, cardiovascular diseases such as heart attacks^17^, artery damage^18^ and strokes^19^. Furthermore, enhancing muscle volume does not necessarily translate into elevated muscle strength^20,21^.

Here we report the identification and first functional characterization of a factor termed myostimulin that is regulated by exercise, augments the regenerative potential and strength of skeletal muscle, and thereby mimics several beneficial effects of exercise without even requiring chronic training or muscle hypertrophy.

## RESULTS

### Exercise elevates muscle *C1orf54*/*C1ORF54* mRNA levels in mice and men

Although exercise is a cornerstone in the treatment of a variety of diseases, its molecular transducers and their complex interactions are not entirely understood ^9,10,14,22^. To discover novel exercise-induced factors, we used a combination of spontaneous wheel running and forced treadmill exercise in mice. This paradigm allowed for preconditioning the animals to physical activity, standardizing the volume of the last exercise bout and tightly controlling the sampling time. By specifically filtering for transcripts regulated by exercise in skeletal muscle, coding for targets previously considered uncharacterized and predicted to be extracellularly located, we found *C1orf54* (Fig. 1a). In fact, we observed that the mRNA levels of *C1orf54* in skeletal muscle increased rapidly in response to exercise and remained elevated for hours after cessation (Fig. 1a). To confirm the relevance of the finding for humans, a publicly available dataset comprising RNA sequencing data from vastus lateralis of young study participants prior and post exercise was re-analyzed^23^. *C1ORF54* mRNA levels markedly increased upon exercise and the elevation was independent of the type of training regimen hence reflecting a general response to exercise (Fig. 1b). Taken together, these data provided the first evidence that *C1orf54*/*C1ORF54* transcription is regulated by muscle contraction in men and mice.

**Fig. 1:**
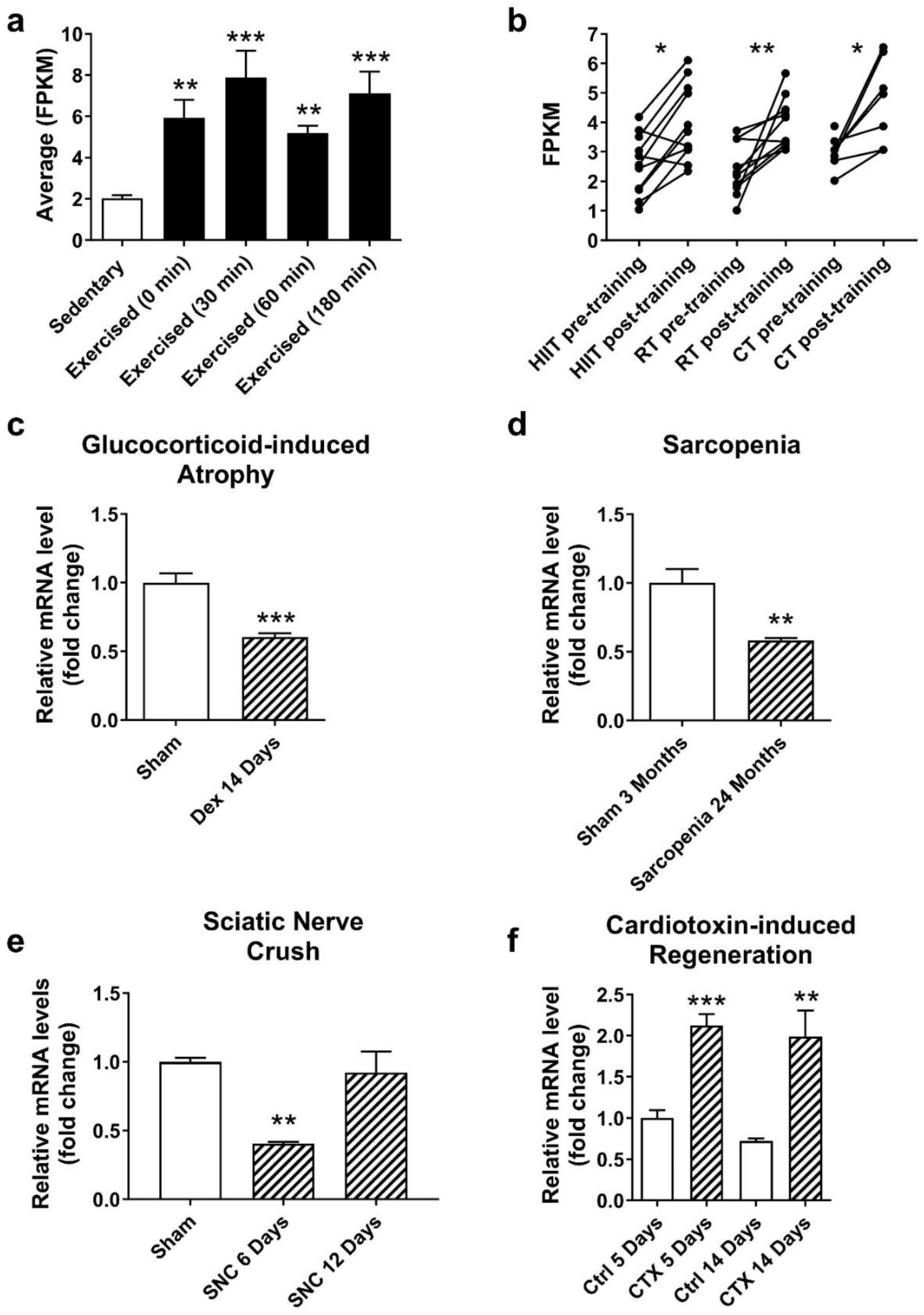
Regulation of *C1orf54/C1ORF54* mRNA levels in trained, atrophic and regenerating skeletal muscle. a) *C1orf54* mRNA levels in mouse quadriceps in response to exercise. Values are expressed as means ± SEM (n=7-8); **: *P* < 0.01, ***: *P* < 0.01 *vs.* sedentary mice by ANOVA and Dunnett post hoc test. FPKM = Fragments per kilobase of transcript per million mapped reads. b) *C1ORF54* mRNA levels in human vastus lateralis from young subjects in response to different exercise paradigms. Individual values pre– and post-training are shown for each subject. n=7-11 participants/group. One subject in the combined exercise training group was excluded from the statistical analysis as no post-training value was available for that subject. All p-values were adjusted using Benjamini and Hochberg multiple testing correction. Statistically significant differences from pre-training are indicated as follows: *p < 0.05, **p<0.01. HIIT = high-intensity aerobic interval training, RT = resistance training, CT = combined exercise training. c and d) Skeletal muscle *C1orf54* mRNA expression under atrophic conditions, specifically in glucocorticoid-induced atrophy (c) and sarcopenia (d). Values are expressed as means ± SEM (n=7-10); **: *P* < 0.01, ***: *P* < 0.01 *vs.* their respective control by unpaired t-test. e and f) *C1orf54* mRNA levels following a sciatic nerve crush (e) and during cardiotoxin-induced regeneration (f). Values are expressed as means ± SEM (n=5-6); **: *P* < 0.01, ***: *P* < 0.01 *vs.* their respective control by unpaired t-test.

### *C1orf54* mRNA levels are reduced in several muscle disorders but elevated during regeneration

Next, we determined mRNA levels of *C1orf54* in mouse models for common human musculoskeletal disorders. Glucocorticoid-induced atrophy represents one of the most frequent drug-induced muscle atrophies^24^, while sarcopenia, characterized by a rapidly progressing decline in muscle mass and function, occurs naturally in mice (≥ 20 months) and humans (≥ 60 years) ^25,26^. Muscle *C1orf54* mRNA levels dropped in both conditions in the mice by 42% and 40%, respectively (Fig. 1c and d). In addition, the levels of *C1orf54* transiently decreased by 60% in mice with a sciatic nerve injury but rebounded during recovery (Fig. 1e). Strikingly, *C1orf54* mRNA levels rapidly rose following cardiotoxin-induced injury (Fig. 1f), a condition characterized by marked satellite cell activation^27^. These data suggest that *C1orf54* expression is reduced in atrophic skeletal muscle and increases in regenerating skeletal muscle (regenerating from exposure to cardiotoxin or exercise).

### Forced overexpression of *C1orf54* evokes a skeletal muscle phenotype

To gain further insights into the potential physiological function of *C1orf54*, we performed overexpression studies with adeno-associated viruses (AAVs). Injection of GFP or bicistronic GFP-C1orf54 AAVs into tibialis anterior infected virtually all muscle fibers (Fig. 2a) and increased *C1orf54* mRNA levels (Fig. 2b) in a dose-dependent manner at doses ranging from 10^10^ to 10^12^ gene copies (GCs). Based on these findings, a dose of 5*10^11^ GCs was selected for further studies (Fig. 2c-f). Mice injected with 5*10^11^ GCs GFP-C1orf54 AAVs showed no overt phenotype one month post infection and had similar muscle weight as control mice injected with GFP AAVs (Fig. 2c). However, transcriptional analyses revealed that overexpression of *C1orf54* significantly dampened the mRNA levels of early drivers of myogenesis, particularly *Myod1* and *myogenin* (Fig. 2d). *Myod1* is a key determinant of satellite cell fate, mainly by committing this cell population to differentiation. Loss of *Myod1* prevents differentiation and results in proliferation as well as improved engraftment and survival when satellite cells are transplanted^28–30^. Hence, the decreased *Myod1* and *myogenin* levels in response to forced constitutive overexpression of *C1orf54* pointed towards a potential involvement of *C1orf54* in regeneration possibly by driving satellite cells to proliferate.

**Fig. 2:**
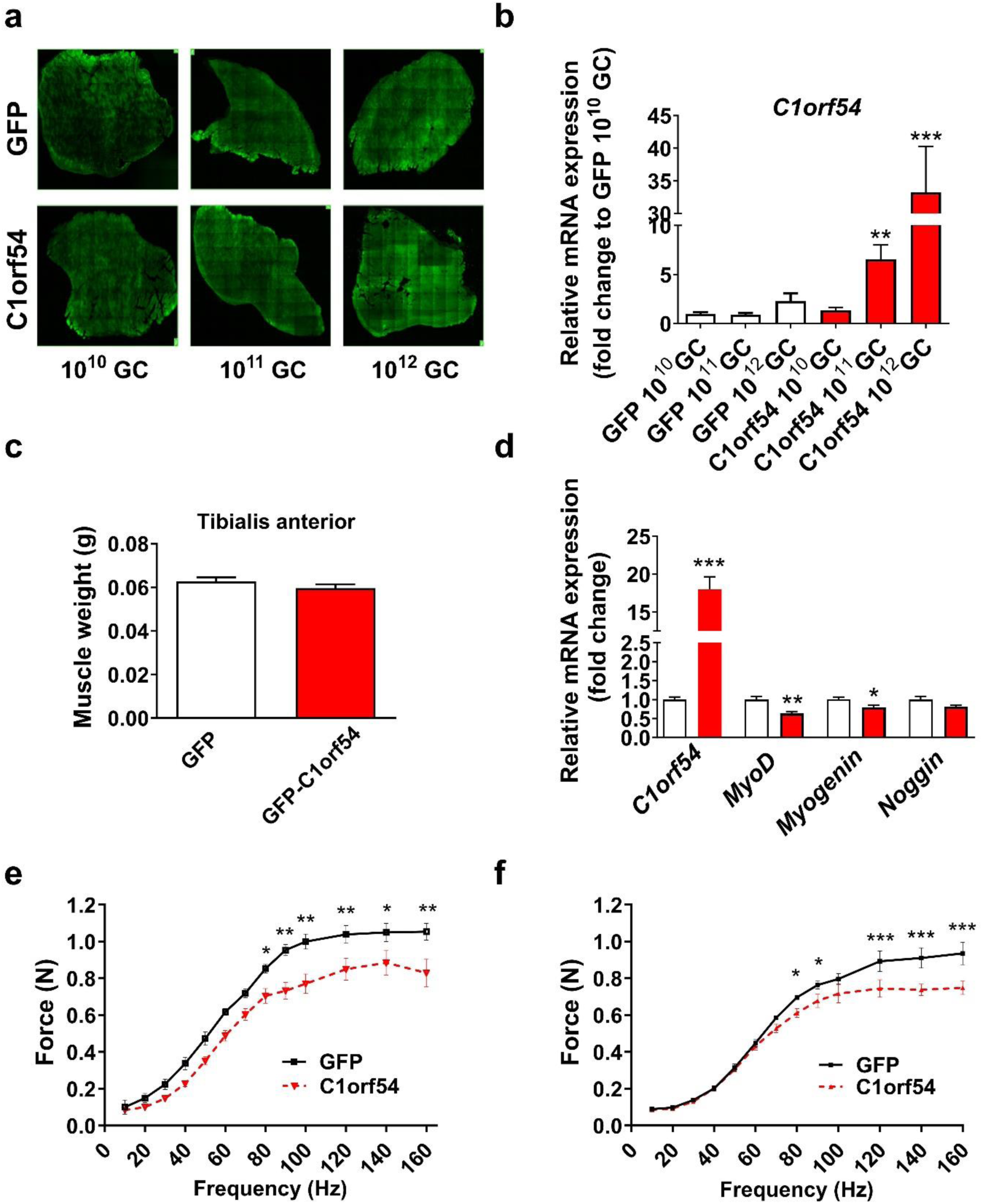
Forced overexpression of *C1orf54* modulates myogenesis and muscle strength. a and b) Imaging of green fluorescence on muscle sections (a) and *C1orf54* mRNA expression (b) one month after intramuscular injection of indicated doses of GFP or bicistronic GFP-C1orf54 AAVs into tibialis anterior. c and d) Tibialis anterior weight (c) and muscle mRNA levels of *C1orf54* and myogenesis markers (d) of mice injected with 5×10^11^ GCs GFP or bicistronic GFP-C1orf54 AAVs into tibialis anterior. e and f) Muscle strength of the injected (e) and the non-injected, contralateral hindlimb leg (f) in response to increasing electrical stimulation. Values are expressed as means + SEM); *: *P* < 0.05, **: *P* < 0.01, ***: *P* < 0.01 *vs.* control by unpaired t-test comparison.

Moreover, constitutive overexpression of *C1orf54* engendered functional adaptations in muscle tissues. Specifically, *C1orf54* considerably decreased muscle strength in the AAV-injected (Fig. 2e) as well as in the non-injected, contralateral leg (Fig. 2f). These data provided evidence that constitutive exposure to high levels of *C1orf54* reduces myogenesis and muscle function, while further suggesting that the protein can be secreted.

### Recombinant C1ORF54 promotes proliferation *in vitro*

Sequence analysis and analytical data indicated that myostimulin is an intrinsically disordered protein (Fig. 3a and Supplementary Fig. 1-10 and Tables 1-3). Applying a recently described algorithm to predict signal peptides and their exact cleavage sites identified the N-terminus as a signal peptide and predicted amino acids 16/17 as cleavage site^31^. We thus inferred that the first sixteen, N-terminal amino acids might be dispensable for activity. Indeed, recombinant C1ORF54 only comprising amino acids 17 to 131 was bioactive and markedly stimulated the proliferation of human myoblasts. Within five days, cells treated with 1000 ng/ml C1ORF54 (aa 17-131) became confluent twice as quickly as those exposed to vehicle (Fig. 3b). We thus named the protein, devoid of amino acids 1-16, myostimulin.

**Fig. 3:**
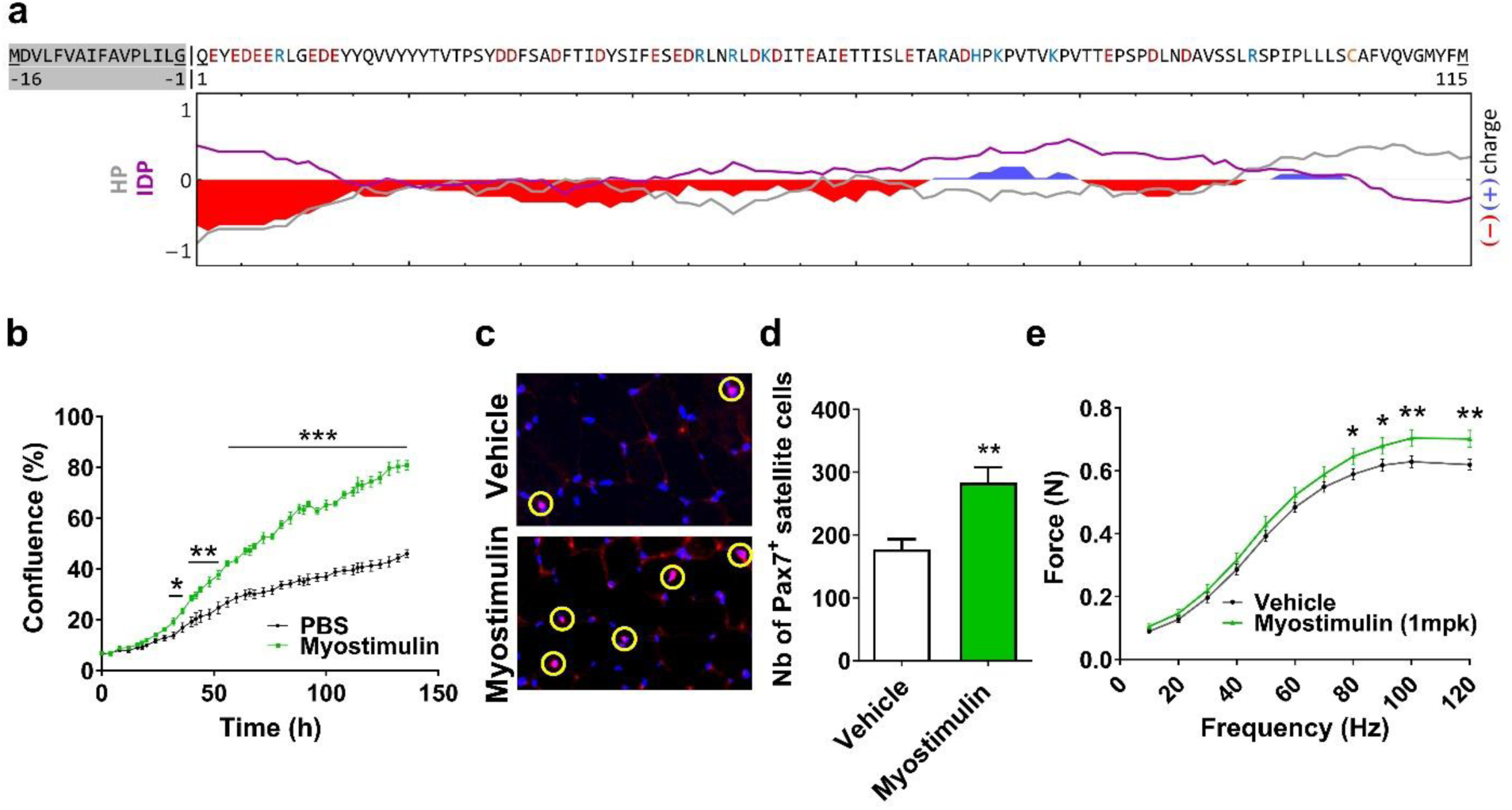
Recombinant, intermittently administered myostimulin increases *in-vitro* and *in-vivo* muscle cell proliferation as well as muscle strength. a) Sequence analysis: Charge, normalized hydropathy (HP) and intrinsic disorder propensity (IDP) plotted for WT myostimulin using a moving window of 15 residues. The C1ORF54 signal peptide is marked in grey. b) Growth curves of primary human myoblasts treated with 1 µg/mL recombinant human myostimulin protein (green squares) and solvent control (black circles) in growth medium containing 1% FBS. Values are expressed as mean ± SEM, n=6 per group. Statistically significant differences are assessed by multiple unpaired t-test comparison and indicated as follows: *p < 0.05, **p < 0.01 and ***p<0.001. c) Staining of muscle sections for nuclei and satellite cells following treatment of mice with myostimulin. Nuclei are shown in blue, satellite cells co-localized with nuclei appear pink and are encircled in yellow. d) Quantification of Pax7 positive satellite cells in mouse gastrocnemius following treatment with vehicle or myostimulin protein. n= 9-10 mice/group. e) Absolute isometric force in mice treated with vehicle or 1mg/kg myostimulin protein. n= 9-10 mice/group. All values are expressed as mean ± SEM. Statistically significant differences are assessed by unpaired t-test for d and e) and indicated as follows: *p < 0.05 and **p < 0.01.

### Myostimulin stimulates satellite cell proliferation and boosts muscle strength *in vivo*

Since exercise evokes a strong, acute stress response, phases of exercise and recovery need to be alternated to allow for adaptive processes. Exercise interventions with insufficient recovery between bouts of exercise lead to over-training which can be associated with declining muscle function^11^. Accordingly, we hypothesized that the impaired myogenesis and reduced muscle function following AAV-mediated constant overexpression were caused by constant exposure to C1ORF54. To avoid continuous exposure to C1ORF54, we dosed mice with 1 mg/kg myostimulin in an intermittent manner (on day 1, 4, 7 and 10). This treatment augmented the number of Pax7 positive satellite cells in gastrocnemius by approximately 66% compared to vehicle treatment (Fig. 3c and d).

We wondered whether the elevated number of Pax7 positive satellite cells was associated with any functional improvements. In fact, myoblast transfer therapy has been previously shown to increase isometric muscle strength in boys suffering from Duchenne muscular dystrophy^32^. Similarly, elevated strength following satellite cell transplantation have been reported in dystrophic mice or in mice with X-linked myotubular myopathy^33,34^. Mice were therefore injected with 1 mg/kg myostimulin protein on day 1, 4 and 7 and then assessed for skeletal muscle force on day 9. At low stimulation frequencies ranging from 10-70 Hz, the evoked force of the hind limb was similar between animals treated with vehicle and myostimulin (Fig. 3e). By contrast, at higher stimulation frequencies ranging from 80-120 Hz and reflecting maximal force generation, mice treated with myostimulin displayed a 13% higher absolute muscle force than vehicle treated animals (Fig. 3e).

### Myostimulin binds to various extracellular receptors

To gain first insights into the potential mode of action of myostimulin, we determined the cellular localization of exogenously administered protein using a myostimulin-azide conjugate and biorthogonal chemistry. In the presence of Dibenzocyclooctyne-Bodipy (DBCO-Bodipy), the azide group in myostimulin reacts with DBCO-Bodipy to become fluorescently labelled. The small azide tag should have a minimal influence on the protein structure and localization. Human myoblasts were seeded in 96 well plates and treated with myostimulin-azide. DBCO-Bodipy was added to localize myostimulin and the labeling revealed a fiber-like structure resembling the extracellular matrix (ECM). To show the extracellular localization of the signal, we used Trypan blue, which is a non-cell-permeable quencher. In the absence of Trypan blue, a green signal was clearly detected, while the presence of the dye quenched the green signal, demonstrating that the azide-tagged myostimulin remained outside the muscle cell (Fig. 4a). To corroborate the co-localization of myostimulin with the ECM, we co-stained with antibodies against different ECM proteins and integrins. A co-localization signal could be observed with fibronectin and integrin β1 (Fig. 4b). Given the extracellular localization, the question arose whether receptors for myostimulin could be identified. To this end, we used the Retrogenix technology enabling screening against a large portion of the human surfaceome, as an unbiased approach. The assay was run on fixed cells that were first incubated with the myostimulin protein. Then, the binding hits were detected by a proprietary myostimulin-specific antibody (IPAL0815) coupled to a fluorescent Alexa Fluor 647 dye (Fig. 4c). The Retrogenix panel yielded 4 plasma membrane receptors and 1 secreted receptor to which myostimulin bound with intensities deemed to be at least weak in both replicates (Fig. 4d red circle). In addition, 16 secreted binding proteins were detected (Supplementary Table 4). A proprietary protein microarray confirmed binding of myostimulin to PLXDC2 (Plexin Domain Containing 2) and the sIL6R (soluble Interleukin-6 Receptor) (Fig. 4d blue circle and Supplementary Table 4). Transcriptomics of Pax7 positive satellite cells freshly isolated from untreated mice revealed that satellite cells express high levels of PLXDC2 and gp130, the membrane-bound coreceptor for sIL6R (Fig. 4d green circle and Supplementary Table 4). Using diverse approaches, we thus demonstrated consistent binding of myostimulin to PLXDC2 and the sIL6R, and that these receptors and coreceptors were highly expressed in muscle precursor cells (Fig. 4d).

**Fig. 4:**
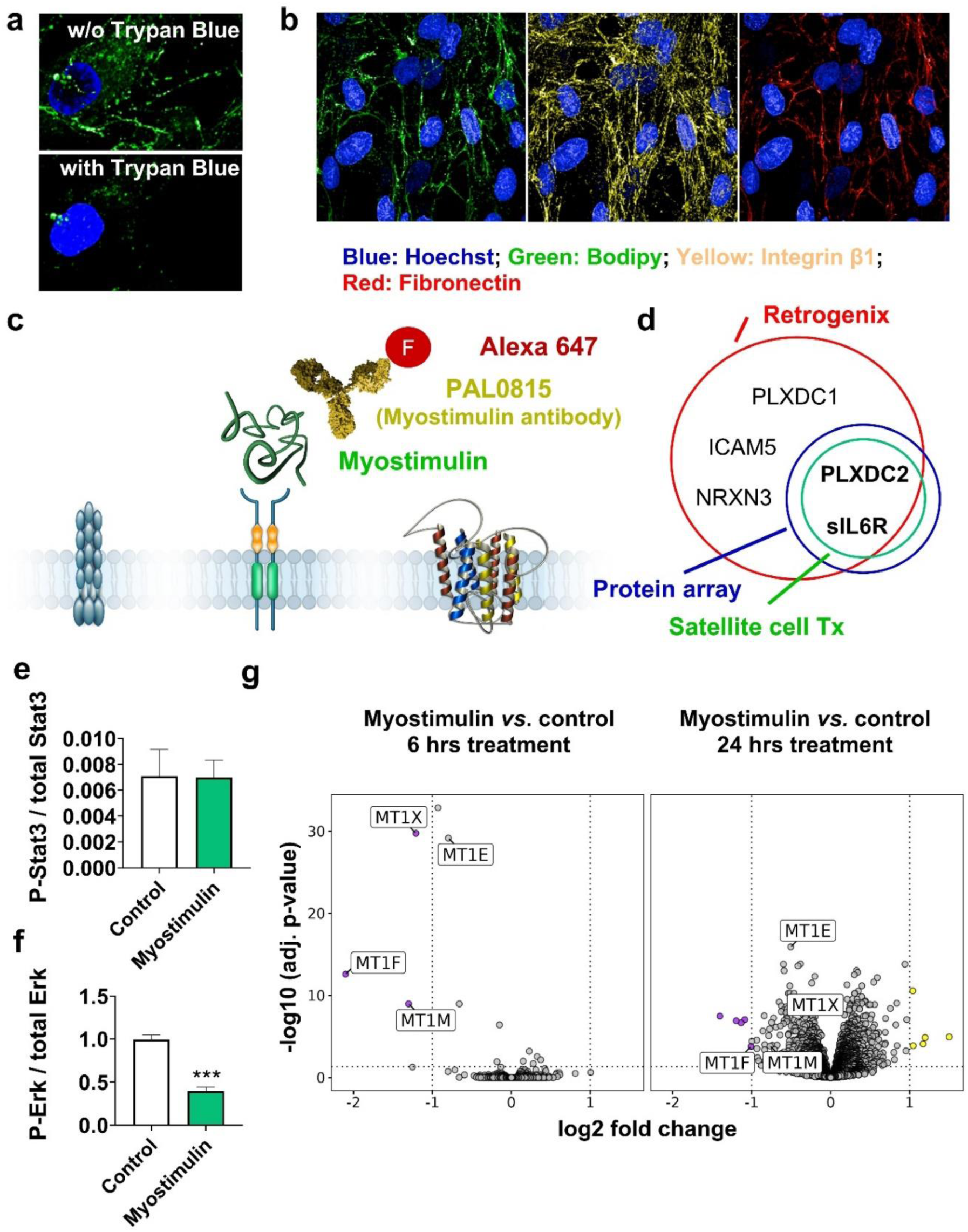
Myostimulin binds diverse receptors and inhibits metallothionein expression. a) Cellular localization of myostimulin. Human myoblasts were treated with myostimulin-azide and then reacted with the green dye DBCO-Bodipy. The disappearance of the green fluorescence after addition of trypan blue, a non-cell-permeable quencher, indicates extracellular localization of myostimulin. b) Myostimulin-azide (green) co-localizes with fibronectin (red) and integrin β1 (yellow) in human myoblasts suggesting ECM localization. c) Schematic of the Retrogenix principle. Fixed cells expressing extracellular receptors are treated with myostimulin. Binding of myostimulin to a specific receptor is detected by a conjugate of an anti-myostatin antibody and an Alexa Fluor 647 fluorophore. d) Receptor hits from the Retrogenix screen (red circle) and additionally showing consistent binding to myostimulin by protein arrays (blue circle). Hits with receptors or coreceptors highly expressed in satellite cells are displayed in the green circle. e and f) STAT3 (e) and ERK (f) phosphorylation in human skeletal muscle cells treated with myostimulin. g) Volcano plots displaying differentially expressed genes between myostimulin-treated human skeletal muscle cells and controls (6– and 24-hours post treatment) as identified by linear regression analysis. Significantly (BH adjusted p-value < 0.05) up– and down-regulated (absolute log2 fold change > 1) genes are colored in yellow and purple, respectively.

While little is known about the further downstream signaling of PLXDC2, it is well established that sIL6R signals via the STAT3 and/or the ERK pathway. When we treated human skeletal muscle cells with myostimulin for 5 minutes, we could not detect any change in the phosphorylation of STAT3 (Fig. 4e). By stark contrast, myostimulin significantly decreased the phosphorylation of Erk (Fig. 4f). Phospho-ERK translocates to the nucleus where it regulates gene transcription. To further elucidate the downstream events, human skeletal muscle cells were treated with myostimulin for 6 and 24 hrs before transcriptomics was performed. Intriguingly, these analyses unveiled a cluster of mRNAs that were consistently downregulated by myostimulin, and which comprised the metallothioneins MT1F, MTFM and MT1X (Fig. 4g).

### Engineered myostimulin with improved biophysical properties and enhanced activity

Myostimulin contains a cysteine residue at position 105 (Fig. 3a) and can therefore form an interchain disulfide bridge. We observed 30-40 % covalent dimers in freshly purified myostimulin and a slow increase in this fraction during long-term storage at 4 °C. In addition to incomplete disulfide formation, other modifications were detected by LC-MS, including acetylation and oxidation, but also +32 Da and +64 Da adducts, that were exclusively found in the dimeric fraction (Fig. 5a). These observations prompted us to engineer a myostimulin variant that was less sensitive to these post-translational modifications (PTMs). To that end we replaced Cys105 by Trp and Phe107 by Leu, based on multiple sequence alignment with myostimulin orthologues from other species (See Supplementary Material). These modifications resulted in a monomeric myostimulin variant, free of the PTMs that were found for the dimeric WT myostimulin (Fig. 5a). Since MT1F is the most myostimulin-responsive metallothionein, we assessed the activity of WT myostimulin and its variant [C105W, F107L] head-to-head using MT1F expression in human skeletal muscle cells as a readout. The variant proved to be more potent than the wild-type protein with an IC_50_ of 102nM for the wild type and 25.2nM for the variant (Fig. 5b). To profile both proteins *in vivo*, we injected 1mg/kg of each protein in an intermittent fashion (day 1. 4, 7, 10 and 13). In a first study the animals were sacrificed two days after the last injection to assess the number of Pax7 positive satellite cells (Fig. 5c). In a second study, muscle strength was evaluated 10 days after the last injection (Fig. 5c). The direct comparison of the two proteins showed that the myostimulin variant tended to be more efficacious in increasing the number of Pax7 positive satellite cells (Fig. 5d) and markedly enhanced muscle strength (Fig. 5e).

**Fig. 5:**
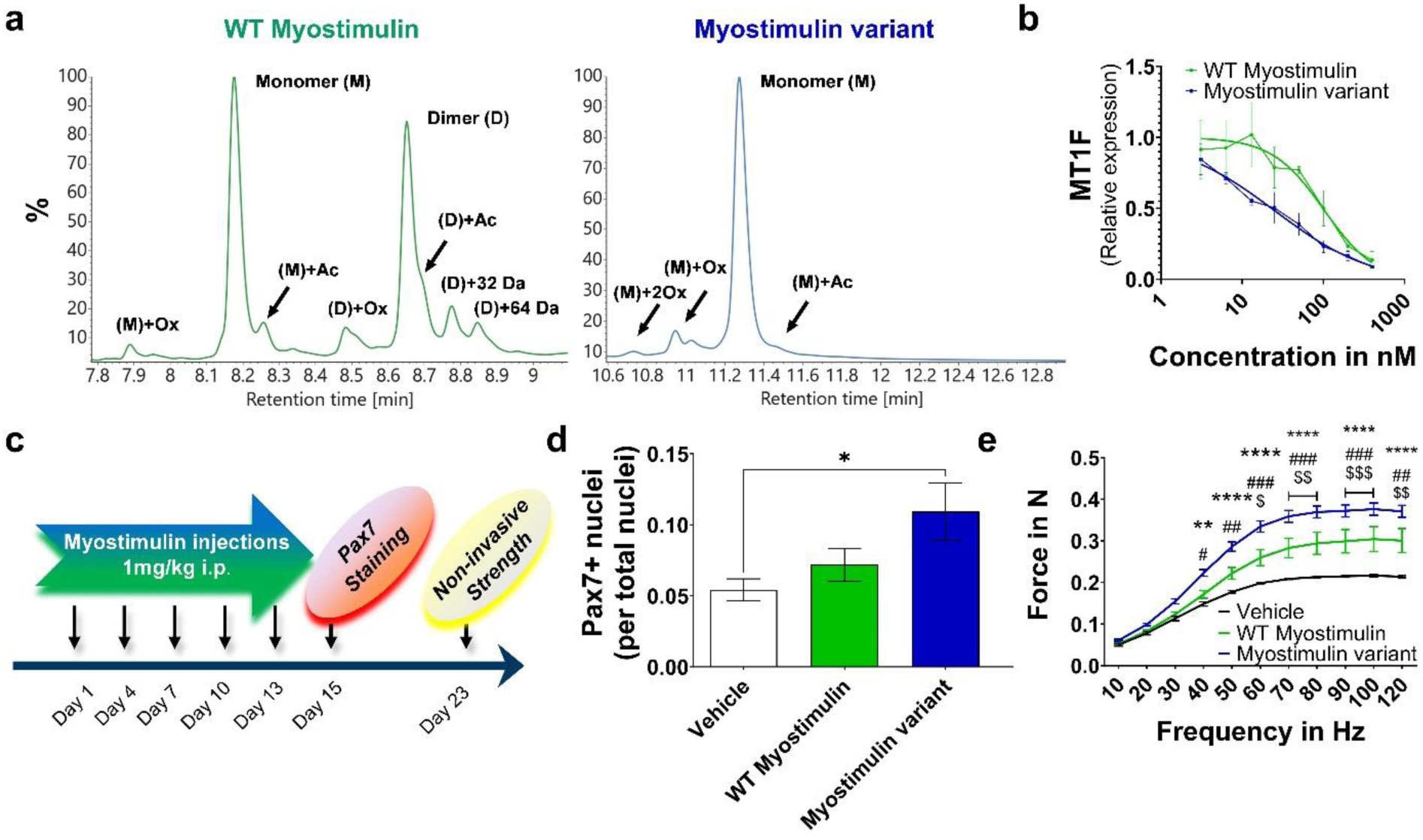
The myostimulin variant [C105W, F107L] features reduced sensitivity to post-translational modifications, and higher *in-vitro* and *in-vivo* activity. a) RP-HPLC-MS revealed partial oxidation (Ox) and acetylation (Ac) of myostimulin. The WT can form covalent dimers (D), while the [C105W, F107L] variant is 100 % monomeric (M) under denaturing conditions. So far uncharacterized PTMs (+32 Da, +64 Da) were detected exclusively for covalent dimers of the WT. b) Relative MT1F mRNA expression in human skeletal muscle cells treated with increasing concentrations of wild type myostimulin or the myostimulin variant (plotted curves include curve fitting). c) Study design for head-to-head profiling of the wild type myostimulin and the myostimulin variant. Animals were sacrificed on day 15 for Pax7 stain or on day 25, 2 days following evaluation of muscle strength d) Quantification of Pax7-positive satellite cells. Values are expressed as means + SEM, n=7-9, Statistical analysis was performed by ANOVA and Holm-Sidak post hoc test; *: *P* < 0.05 as indicated. e) Isometric muscle strength. Values are expressed as means + SEM, n=5-10. Statistically significant differences are assessed by 2 factor ANOVA with Holm-Sidak test comparisons and indicated as follows: *, Vehicle *vs.* myostimulin variant; # WT myostimulin *vs.* myostimulin variant; $, vehicle *vs.* WT myostimulin. One symbol p < 0.05, two symbols p < 0.01, three symbols p < 0.001 and 4 symbols p<0.0001.

### Engineered myostimulin counters the loss of strength following peripheral nerve injury

Since damage of the sciatic nerve reduces the mRNA levels of *C1orf54* (Fig. 1e) and recombinant myostimulin increases muscle function in healthy mice (Fig. 3a and 5e), we sought to determine whether exogenous administration of the myostimulin variant alters muscle function in this injury setting. To this end, we first injured the sciatic nerve to reduce muscle strength followed by treatment with the myostimulin variant. Changes in muscle strength were monitored repeatedly (Fig. 6a).

**Fig. 6:**
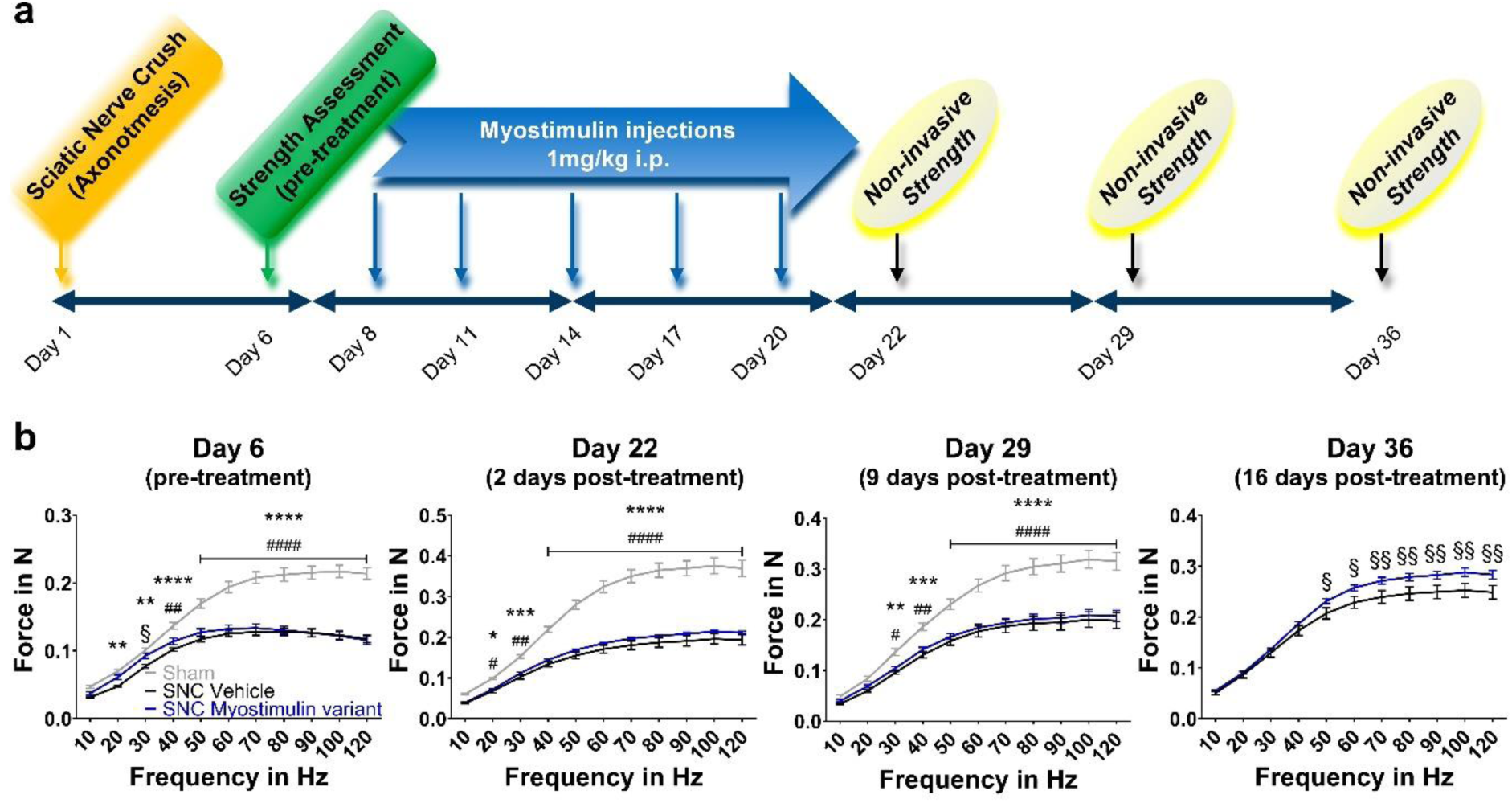
The myostimulin variant [C105W, F107L] alleviates the decline in muscle strength induced by peripheral nerve injury. a) Study design. Axonotmesis was induced by a sciatic nerve crush. Muscle strength was assessed 6 days later, and animals randomized into a vehicle and a myostimulin variant treatment group. A sham group was included to determine the initial degree of muscle dysfunction by the nerve crush. All animals received 5 injections of the myostimulin variant. Following the treatment, the development of muscle strength was monitored for two additional weeks. Of note, the sham group was included in the first measurements, but the measurement of this group was abandoned in follow-up assessments. b) Isometric strength of the operated-leg 6 days after the nerve injury and following treatment with either vehicle or 1mg/kg of the myostimulin variant on day 22, 29 and 36, which correspond to days 2, 9 and 16 post treatment. respectively. n= 8-12 mice/group. All values are expressed as mean ± SEM. Statistically significant differences are assessed by 2 factor ANOVA with Holm-Sidak test comparisons and indicated as follows: *, sham *vs*. sciatic nerve crush (SNC) treated with vehicle; # sham *vs.* SNC treated with the myostimulin variant; §, SNC treated with vehicle *vs*. SNC treated with the myostimulin variant. One symbol p < 0.05, two symbols p < 0.01, three symbols p < 0.001 and 4 symbols p<0.0001.

Damaging the sciatic nerve led to a dramatic decrease in muscle strength within 5 days (Fig. 6b). On day 6 mice which had experienced a loss in muscle function exceeding 0.15 N were randomized into two groups. One group was then treated therapeutically with intermittent injections of 1mg/kg myostimulin or vehicle for 12 days. Two days after treatment completion, the strength curves started to segregate with the difference reaching statistical significance 16 days post-treatment (Fig. 2b). These data provide a preclinical proof of concept that the myostimulin variant can expedite the recovery of muscle strength in a motor nerve injury setting and has potential for conditions with impaired skeletal muscle function.

## DISCUSSION

The burden of musculoskeletal diseases and injuries has markedly increased over the past several decades and is projected to grow substantially as the world population ages and continues the pervasive sedentary lifestyle^1,35^. Currently, musculoskeletal diseases are the leading contributor to years lived with disability worldwide^1^. Consequently, the development of drugs that restore and enhance the function of skeletal muscle in a safe and timely manner will take on greater importance. Here we identified *C1orf54/C1ORF54* as an endogenous factor that increases in skeletal muscle after exercise and during regeneration and decreases in atrophying muscle (Fig. 1a-f). Removing 16 amino acids from the N-terminus of the C1ORF54 protein generates a protein that we named myostimulin. We provide evidence that myostimulin is a bioactive protein that stimulates the proliferation of muscle precursor cells *in vitro*, drives satellite cell expansion *in vivo*, and increases muscle strength (Fig. 3b-e, Fig. 5b, d and e, and Fig. 6b). Importantly, an improved myostimulin variant even maintains its strength-enhancing capacity in diseased muscle and thereby accelerates the recovery of muscle function from axonotmesis (Fig. 6b). Taken together, these data position myostimulin as a novel player in skeletal muscle physiology, a potential biomarker of the health status of muscle, and a promising new target for an innovative pharmacological therapy to counter muscle weakness and restore functional autonomy.

An important aspect of our findings is that exogenous myostimulin is sufficient to stimulate the proliferation of muscle-resident satellite cells *in vivo* (Fig. 3c and d and Fig 5d). Satellite cells maintain and regenerate myofibers by a tightly regulated process involving activation of quiescent satellite cell populations, subsequent proliferation to amplify the cell pool, and generation of new differentiated myofibers or repair of existing, damaged ones^36,37^. Several lines of evidence show that increasing the number of muscle precursor cells can increase muscle strength. In boys with Duchenne muscular dystrophy transplantation of myoblasts has been shown to improve isometric muscle strength^32^. Similarly, satellite cells expanded *in vitro* enhanced muscle strength following transplantation into mice with muscle injuries^38^. However, current approaches focus on expansion of satellite cells *in vitro* and subsequent transplantation, which can lead to a low number of viable cells and the rejection of transplanted cells by the recipient. A major advantage of myostimulin consists of its direct effect on satellite cell proliferation *in vivo* which reduces such risks.

Of note, intermittent dosing with myostimulin rapidly improves maximal isometric muscle strength (Fig. 3e, Fig. 5e and Fig. 6b). Increased muscle strength is a hallmark of long-term, heavy resistance training. A meta-analysis reported that at least 4 weeks of heavy resistance training is required to achieve about 18.4 % higher strength in humans^39^. In rodents, improving muscle strength proved to be much more challenging. Few approaches have been shown to elevate isometric muscle force with resistance exercise paradigms in rodent models and the change in force is invariably associated with higher muscle mass^40–42^. Given the difficulty in evoking characteristic effects of resistance training in a standard pre-clinical model, the finding that myostimulin enhances strength in mice, particularly in the absence of muscle hypertrophy, is remarkable. In humans, strength measured isometrically correlates with mobility parameters such as chair rise^43^ and gait^44^, and isometric force/body weight ratio is a reliable measure for modeling performance of ambulatory tasks^45^. Treatment with myostimulin might thus represent a modality to potentially increase functional capacity (i.e., sit-to-stand, mobility) and to enhance the effect of rehabilitation and exercise programs; thereby improving the quality of life.

Being an intrinsically disordered protein, myostimulin can assume different confirmations and might interact with various proteins, including cell surface receptors. We show that myostimulin binds to PLXDC2 and the sIL6R (Fig 4d). PLXDC2 is a transmembrane protein and a cell surface receptor for Pigment Epithelium-Derived Factor (PEDF), a growth factor that has been shown to promote skeletal muscle regeneration via mitogenic effects on muscle precursor cell^46,47^. By contrast, sIL6R represents a soluble, extracellular receptor. This receptor transactivates cells by binding to gp130, a transmembrane receptor. Interestingly, we found that gp130 is particularly highly expressed in muscle precursor cells (Supplementary Table 4). The pathway signals via STAT3 and/or ERK and can increase the expression of metallothioneins^48^. Our data establishing binding of myostimulin to sIL6R, reducing Erk phosphorylation and MT1 expression (Fig. 4d-g), suggest a consistent modulatory effect of myostimulin on this pathway. The finding that myostimulin ultimately inhibits metallothionein expression is in accordance with our previous work demonstrating that blockade of metallothioneins in mice significantly increases muscle strength^49^. Evidently, as an intrinsically disordered protein, myostimulin may mediate its effects via different mode of actions and further investigations will be required to elucidate whether additional pathways are involved. Nonetheless, our data provide first evidence of an involvement of the sIL6R/gp130 coreceptors, reduced downstream Erk phosphorylation and decreased metallothionein expression in promoting a resistance-trained phenotype with myostimulin treatment.

The therapeutic potential of myostimulin is intriguing. Exogenously administered myostimulin exhibits dual beneficial effects on skeletal muscle, and it presumably could be prepared and delivered as an injectable polypeptide for clinical use. An obstacle to the development of myostimulin as a therapeutic is its susceptibility to post-translational modifications. The myostimulin variant we developed does not undergo covalent dimerization and is free of the uncharacterized post-translational modifications we observed for the wild-type protein. The variant also displays enhanced biological activity (Fig. 5a-e). Importantly, our data demonstrate that short, low dose treatments with the myostimulin variant ameliorated skeletal muscle function in mice with severe functional limitations due to a peripheral nerve injury, highlighting the efficacy of the myostimulin variant in the preclinical model (Fig. 6b).

We report the identification of a novel protein we named myostimulin and the development of a viable variant with salutary benefits to skeletal muscle function. The worldwide increase in the prevalence and burden of musculoskeletal disorders and associated functional decline create a large unmet need for interventions, like partial exercise-mimetics, with effects on skeletal muscle function. While our findings indicate a potential novel therapeutic avenue to treat skeletal muscle disorders, myostimulin also may hold promise for targeting additional clinical indications. Indeed, the wealth of public data linking exercise numerous health benefits in various other diseases suggests that myostimulin might have disease-modifying effects beyond musculoskeletal diseases and thus potential for broader therapeutic applications.

## ONLINE CONTENT

Any methods, supplementary information, acknowledgements, details of author contributions and competing interests; and statements of data are available online as supplementary information.

## MATERIAL AND METHODS

### Analysis of human exercise data

Published transcriptomics data (GSE97084) from vastus lateralis biopsies from humans undergoing various types of exercise were re-analyzed and counts reported as FPKM (Fragment Per Kilobase per Million)^23^.

### Animals

All animal studies described were performed according to the official regulations effective in the Canton of Basel-City, Switzerland. Adult male C57BL/6 and B6-Pax7tm1(IRES-EGFP) Npa mice were bred at Charles River Laboratories. After arrival at our research facility the mice were acclimatized for seven days. They were housed at 25°C with a 12:12 h light-dark cycle and fed a standard laboratory diet containing 18.2% protein and 3.0% fat with an energy content of 15.8 MJ/kg (Nafag, product # 3890, Kliba, Basel, Switzerland). Food and water were provided *ad libitum*.

### Exercise study in mice

For exercise studies, C57BL/6 mice were granted continuous access to running discs (Med Associates Inc) for seven days. On day 8, the animals were additionally subjected to a single bout of exercise on a treadmill (Panlab, Harvard Apparatus). In brief, mice were adapted to the treadmill for 10 min at a speed of 14m/min. Thereafter, the speed was increased gradually by 2m/min every 4 minutes until it reached 26m/min. The final speed of 26m/min was maintained for 4 min. Mice were then euthanized at specific time points, i.e. immediately, 30-, 60-or 180-minutes post-treadmill exercise.

### Musculoskeletal disease models

To simulate glucocorticoid-induced muscle atrophy, mice received 1.25 mg/kg/d dexamethasone-21-phosphate p.o for 14 days^49^. To study sarcopenic skeletal muscle, mice were aged up to 24 months and compared to adult 3 months old animals. Peripheral nerve injury was mimicked by exposing the sciatic nerve and squeezing the nerve with a forceps. Sham-operated animals were used as controls. For studying muscle regeneration, mice received a single injection of 25µl cardiotoxin (10µM) into their tibialis anterior muscle. Following the respective intervention, the animals were euthanized, and skeletal muscle tissue was collected for post-mortem analyses.

### Gene expression analyses of muscle tissue

For gene expression profiling from muscle tissue, total RNA was extracted from muscles using TRIzol reagent (Invitrogen) according to manufacturer’s instructions and processed as previously described^49^. RT-PCR results were expressed as fold changes over controls and RNA-seq results were expressed as FPKM (Fragments Per Kilobase Million).

### In-vivo target validation with AAVs

For studies involving AAVs, adult C57BL/6 mice received a single intramuscular injection of AAV expressing GFP or bicistronic GFP-mouse C1orf54 (Vectorbiolabs) at the dose specified. Mice were euthanized one-month post-injection.

### Muscle strength

The strength of the hind leg was assessed noninvasively as previously described^49,50^.

### Proliferation of primary human skeletal muscle cells in vitro

To measure proliferation in vitro, primary human skeletal muscle cells (myoblasts) from Cook MyoSite, Pittsburgh, PA, USA were cultured in growth medium (GM) containing MyoTonic serum-free growth medium (COOK MK-2222), growth supplements (COOK MK-8888), 10% fetal bovine serum superior (Millipore, S0613, lot 15H095) and 50 µg/mL gentamycin (Gibco, 15710-049). After overnight incubation, cells were incubated with GM without FBS for 2.5 h, followed by medium change into GM containing 1% FBS for 2.5 h. Subsequently, cells were treated with 1000 ng/mL recombinant human C1ORF54 protein aa 17-131 (Mybiosource, MBS 1284901 lot 03095), or PBS control. Cell growth as percent (%) confluence was measured by life cell imaging using an IncuCyte FL Live-Cell analysis system (Essen BioScience, Ltd.).

### Recombinant expression and purification of myostimulin

The DNA sequences encoding WT myostimulin (human C1orf54, residues 17—131) and its variant [C105W, F107L] were optimized for expression in E. coli using an unpublished, in-house algorithm, synthesized at GeneArt and cloned into pET-26b expression vectors. To facilitate complete post-translational removal the start methionine, a Ser codon was added at position 2. In addition, His6-NproEDDIE-tagged (Supplementary Discussion) constructs of the WT and the variant were generated and cloned into pET-30. E. coli BL21 Gold (DE3) cells were transformed with the respective plasmid. Protein expression was carried out in Terrific Broth under batch fermentation conditions at 37 °C in a DASGIP® bioreactor (Eppendorf). Cells were harvested 3 hours post induction and frozen immediately. Inclusion bodies (IB’s) were isolated and washed as described by Rudolph et al.,^51^.

IB’s of the non-His-NproEDDIE-tagged myostimulin were solubilized in 6 M urea, 20 mM Tris pH 8.0, 5 mM EDTA, 0.2 M sodium sulfite, 5 mM DTT. The solution was filtered and purified over an 80 mL Poros R1 (20 µm) reversed-phase column, placed in a TL-105-D column oven (Timberline Instruments) set to 50 °C, connected to an ÄKTA pure 25 system (Cytiva) and equilibrated with 20 mM ammonium bicarbonate. Elution was performed by applying a linear gradient from 0 to 70 % ethanol in 20 mM ammonium bicarbonate over 8 column volumes at 8 ml/min. Up to 30 mg of solubilized protein were injected per run. Fractions containing the protein of interest were pooled and dialyzed against PBS, pH 7.3 supplemented with 1 mM EDTA and 3 mM Met.

His-NproEDDIE-tagged myostimulin IB’s were solubilized in 6M guanidium chloride, 50 mM sodium phosphate pH 8, 1 mM EDTA, 5 mM DTT, loaded on a 5 mL HisTrap Excel column (Cytiva) eluted by pH shift to 4.5. Autocleavage of the NproEDDIE fusion tag was triggered by rapid 1:50 dilution into 1 M Tris/HCl, 0.25 M sucrose, 2 mM EDTA, and 15 mM thioglycerol, pH 7.3 followed by overnight incubation at 4 °C. Uncleaved fusion protein and cleaved His-NproEDDIE were removed using a 5 ml HisTrap Excel column in flow-through mode in 6 M urea. The flow-through fraction was subjected to RP-HPLC as described above for untagged myostimulin. Fractions containing the protein of interest were pooled and dialyzed against PBS, pH 7.3 supplemented with 1 mM EDTA and 3 mM Met.

Purified myostimulin was stored at 4 °C and protected from light. (See Supplementary Discussion.)

### Treatment of mice with recombinant myostimulin

For animal studies involving recombinant proteins, C57BL/6 mice received intraperitoneal injections of vehicle (PBS) or 1mg/kg body weight of recombinant myostimulin formulated in vehicle and administered intermittently with three-day intervals. Mice were euthanized at the time-point indicated.

### Histology and Pax7 staining

Histology for satellite cell quantification was performed on mice that received intermittent injections of 1mg/kg recombinant myostimulin (day 1, 4, 7 and 10 for the data displayed under Fig. 3c and d. and on day 1, 4, 7, 10 and 13 for the data displayed under Fig. 5d). For the data shown in Fig 3, animals were euthanized two days after the last dosing and the gastrocnemius muscle was rapidly dissected out, embedded in OCT (Tissue-Tek) and frozen in liquid nitrogen-cooled isopentane. Frozen muscles were cut into 10 μm cross-sections, which were dried at room temperature for 2 hours and fixed with 3% paraformaldehyde for 10 min. Sections were then blocked with Vector M.O.M blocking reagent (Vector Laboratories, Cat. #MKB-2213, diluted 1/100) and 10% goat serum in PBTX (PBS containing 0.2% Triton X100) for 30 minutes. To detect satellite cells, the sections were incubated overnight with a primary antibody against Pax7 (Developmental Studies Hybridoma Bank (DSHB)) in blocking buffer at 4°C. After several washes with PBTX, the sections were incubated at room temperature with a bridging antibody (rat anti-mouse IgG, Serotec, Cat. #MCA336, diluted 1/500) in 10% goat serum in PBTX for 3 hours. Samples were subsequently washed with PBTX and stained with a fluorescently labeled secondary antibody (Alexa 594 goat anti-rat, Invitrogen, Cat. #A11007, diluted 1/200) at room temperature for 1 hour. After washing with PBTX, samples were mounted with ProLong Gold antifade reagent with DAPI (Invitrogen). Cells positive for both Pax7 and DAPI were counted. The data shown in Fig 5 were generated using B6-Pax7tm1(IRES-EGFP) Npa mice and an anti-GFP antibody (ab6556, Abcam) was used for the detection of satellite cells.

### Image analysis for Pax7 positive cells

Image analysis was performed as described previously^50^. In brief, glass slides were scanned with a Philips Ultra-Fast Scanner (Philips AG, Horgen, Switzerland) at 400x magnification resulting in image pixel dimensions of 0.25 µm × 0.25 µm. For the quantitative assessment of Pax7 positive nuclei, the HALO image analysis platform (Indica Labs, Albuquerque, NM, United States) was used. The image analysis was performed in two steps: first detecting the regions of interest (ROIs) in the tissue (excluding areas that may result in artifacts, e.g., tissue folds) and subsequently detecting nuclei positive and negative for Pax7. For the ROI detection a machine learning based classifier (DenseNet, HALO AI) has been devised and trained to recognize three different classes: muscle tissue, exclusion areas and whitespace. In a second step, nuclei positive and negative for Pax7 have been detected in the previously defined region of muscle tissue using the HALO module Multiplex IHC v3.0.1.

### Bio-orthogonal reaction and co-localization of myostimulin with ECM components

Human myoblasts were seeded in 96 well plates at a density of 15000 cells per well and incubated overnight at 37°C and 5% CO_2_. Myostimulin-azide was added at 100 nM for 15 minutes at room temperature.

Cells were then treated with a solution of 0.3 µM Bodipy-FL-PEG4-DBCO (Jena Bioscience) in DPBS with Ca^2+^ and Mg^2+^ for 45 minutes at room temperature. A volume of 100 µL solution was removed from each well and 100 µL Hoechst dye (Invitrogen) at a final dilution 1:10000 in DPBS with Ca^2+^ and Mg^2+^ was added for 30 minutes. Cells were imaged with a CV7000 at 60x magnification.

To half of the wells in the plate a solution of 50 µl trypan blue at a final concentration of 0.04% was added manually and incubated for 20 minutes at room temperature in the dark. Afterwards, cells were washed manually three times with DPBS with Ca^2+^ and Mg^2+^ and fixed with paraformaldehyde (4% v/v in DPBS). Plates were imaged with a CV7000 high throughput microscope with a 60x magnification objective.

For immunofluorescent labeling, fixed samples were treated with 100 µl of blocking buffer (10% FCS, 0.1 % TritonX100 in PBS) at RT for 30 minutes. 50 µl of volume was removed from wells and 50 µl primary antibody solutions in blocking buffer were added to each well manually for 1h at room temperature. The following antibodies were used: Rabbit polyclonal anti-fibronectin (Abcam, ab2413); rabbit polyclonal anti-collagen type 1 (Rockland, #600-401-103-0.5); rabbit polyclonal anti-laminin (Abcam, ab11575); mouse monoclonal activated anti-integrin β1 (Sigma, #MAB2079Z); mouse monoclonal anti-fibronectin (Abcam, ab281574). Cells were washed three times with 100 µl DPBS with Ca^2+^ and Mg^2+^ and 100 µl secondary antibody added and incubated for one hour: Alexa Fluor 647 goat-anti rabbit antibody (Invitrogen, A21244) and Alexa Fluor 555 goat-anti mouse antibody (Invitrogen, A21422). Cells were washed again with DPBS containing Ca^2+^ and Mg^2+^ and plate measured with a CV7000 high throughput microscope with a 60x magnification objective using the following settings: Hoechst: BP 445/45; Bodipy: BP 525/50; Alexa Fluor 647: BP 676/29; Alexa Fluor 555: BP 617/67.

### Retrogenix

A Retrogenix human cell-based protein array screen was performed at Charles River Discovery Research Services UK. Briefly, 3µg/ml of WT myostimulin was assessed for binding against fixed HEK-293 cells overexpressing individually 4064 human plasma membrane proteins and 2037 secreted proteins (either tethered or non-tethered to the plasma membrane). Additionally, the screening included fixed HEK-293 cells co-expressing 2 proteins known to form heterodimers (396 heterodimers). Binding was detected using a proprietary anti-myostimulin antibody (IPAL0815) labeled with Alexa Fluor 647. ZsGreen1, a green fluorescent protein co-expressed with each target, was used to confirm successful transfection and ensure that a minimum threshold of transfection was achieved. Hits (duplicate spots: 1 and 2 in Supplementary Table 4) were identified by analyzing Alexa Fluor 647 and ZsGreen1 fluorescence on ImageQuant.

### Protein array

To independently identify human proteins interacting with myostimulin, profiling studies were conducted using a proprietary protein microarray platform. This platform comprised a protein library of 6822 proteins representing 2589 unique genes from the human extracellular proteome, including secreted or extracellular domains of single-pass transmembrane proteins. The protein library was printed with a non-contact printer (Marathon, Arrayjet) on ultra-thin nitrocellulose-coated glass slides (PATH slides, Grace Bio-Labs). The incubation process involved multiple steps, including washing, blocking, and incubation with the detection antibody, all performed at a constant temperature of 23°C on the HS 4800 Pro Hybridization Station (Tecan). A myostimulin-Fc recombinant protein was incubated on the protein microarrays at concentrations of 100 nM and 1 µM. Data were pooled in the final analysis. Detection was achieved using an Alexa Fluor647-conjugated goat anti-human-IgG-Fc fragment-specific antibody (Jackson IR Cat#109-605-098). After a drying step at 30°C under a flow of nitrogen, the microarrays were scanned using a GenePix Pro 4200A scanner (Molecular Devices). Data analysis was performed using GenePix Pro v6.1 (Molecular Devices) and Protein Chip Reporter (Novartis). For those proteins identified in the Retrogenix assay (apart from protein NPPB, which was not covered by the protein arrays), the mean fluorescence intensity obtained in the protein arrays was normalized for each of the spotted proteins and expressed as a percentage of the reference spots (human IgG, Jackson IR Cat#009-000-003).

### Sorting of satellite cells and AmpliSeq-targeted transcriptomics

B6-Pax7tm1(IRES-EGFP) Npa mice were euthanized, and muscle-resident cells were isolated from hind limb legs (pooled from several mice) with the Skeletal Muscle Dissociation Kit (Miltenyi Biotec #130-098-305). The pellet of cells has been resuspended in FACS buffer.

EGFP-positive satellite cells were isolated from other muscle-resident cells by preparative flow cytometry cell sorting (Aria Fusion, Becton Dickinson, equipped with BD FACSDiva Software), using the 70 µm nozzle at 70 psi pressure, and 1× BioSure Preservative-Free Sheath Solution (Concentrate, catalog no. 1027). The temperature of the samples and of the collecting tubes was set at 4 °C and cells were sorted using the purity mode and a low event rate to maintain a high recovery yield and limit dead volume loss in rare sample. Cells were sorted into 1.5 ml Eppendorf tubes containing 350 µl of Qiagen RLT Lysis buffer. Total RNA was extracted from FACS-sorted cells using the Quiagen RNeasy micro kit (74004) following the manufacturer’s protocol. RNA concentration and purity were assessed using the Agilent high sensitivity RNA ScreenTape assay on the Agilent TapeStation. RNA libraries were prepared using the Ion AmpliSeq Transcriptome Mouse Gene Expression Kit (A36553; Thermo Fisher Scientific). Briefly, 2.5 ng of total RNA per sample was reverse transcribed into cDNA using the Ion AmpliSeq Library Kit Plus. The cDNA was subsequently amplified in 14 PCR cycles using the Ion AmpliSeq Transcriptome Mouse Gene Expression core panel (23930 targets). Following amplification, primer sequences were partially digested, and the amplicons were ligated to IonCode barcode adapters (A29751; Thermo Fisher Scientific) to enable multiplexing.

The prepared libraries were quantified using the Ion Library TaqMan Quantitation Kit (4468802; Thermo Fisher Scientific) and diluted to the appropriate concentration for template preparation in 96-well reaction plates (4306737; Thermo Fisher Scientific). Template preparation and chip loading were performed using the Ion Chef System using the Ion 550 Kit-Chef (A34541; Thermo Fisher Scientific). Emulsion PCR was conducted to clonally amplify the library fragments on Ion Sphere Particles (ISP), and the ISPs were enriched and loaded onto Ion 550 Chips (A34538; Thermo Fisher Scientific). Sequencing was performed on the Ion S5 System (Thermo Fisher Scientific).

For data analyses, raw sequencing data were processed using the Torrent Suite Software v.5.18 (Thermo Fisher Scientific). Base calling, adapter trimming, quality filtering, and alignment to the mouse reference genome (mm10) were conducted using the default parameters of Torrent Suite. Gene expression levels were quantified using the ampliSeqRNA v5.16 plug-in available in the Torrent Suite.

### Signaling pathway analyses, transcriptomics and RT-PCR in human myoblasts

Primary human muscle cells (hSkMDC19, P201076) passage 7 were thawed and cultured for 5 days in growth medium (MK-2288, COOK Myosite) containing 20% fetal bovine serum (S0615, Millipore), 50 µg/mL gentamycin (15750, ThermoFisher) and human recombinant insulin (5-79F00-G, Amimed) according to standard procedures. Cells (100’000 per well in 2 mL) were seeded into 6WP (Costar 3516) for signaling pathway or 48WP for gene expression analyses in complete growth medium. The next day, cells were starved for 2h by cultivating in starvation medium (growth medium without FBS and gentamycin) containing 25 mM HEPES pH7.4.

For signal pathway analyses, cells were treated for 15 min by medium change with solvent control (PBS) or 200 nM myostimulin. Proteins were pre-incubated for 15 min before treatments. Then, cells were washed with 1 mL PBS (Cat no 14190, GIBCO) followed by addition of 100 µL RIPA buffer (Sigma R0278) containing 5 µM EDTA and 1x Halt protease and phosphatase inhibitor cocktail (ThermoFisher 78440) for 20 min on ice. Cells were scraped from the plate and the lysate was centrifuged for 15min with 13,000 rpm at 4°C. Total protein concentrations were determined using BCA protein assay (ThermoFisher #23227) and aliquots were frozen at –80°C. P-STAT3, STAT3, P-ERK1/2 (T202/Y204) and ERK1/2 were analyzed by JESS Western blots using anti-pSTAT3 (Cell Signaling #9131L), anti-STAT3 (Cell Signaling #4904s), anti-pERK1/2 (T202/Y204) (#9101L Cell Signaling) and anti-ERK1/2 (#9102L Cell Signaling) antibodies.

For RNA-seq analysis, cells were treated for 6h or 24h in triplicates by medium change with solvent control (PBS) or 200 nM myostimulin. For RNA extractions, cell medium was removed by pipetting and centrifuging followed by freezing the plates with cells at –80°C. Subsequently, RNA extractions (Qiagen,74106) were performed.

For RT-PCR experiments to measure MT1F expression, cells were treated for 24h in duplicates by medium change with solvent control (PBS), WT myostimulin or the myostimulin variant at 400 nM and 2-fold serial dilutions as indicated below in the figures. For RNA extractions, cell medium was removed by pipetting and centrifuging followed by freezing the plates with cells at –80°C. Subsequent RNA extractions were performed using RNeasy micro kit (Qiagen) and RT-qPCR of MT1F using corresponding TaqMan assays. Expression of the house keeping gene TBP was used for normalization.

### Analytical Reversed-Phase Chromatography and peak characterization by mass spectrometry

Reversed-phase mass spectrometry experiments were performed by injecting 1µg of sample on Waters Acquity H-class UPLC connected to a Waters Vion IMS QToF mass spectrometer. Protein species separation was achieved by applying a gradient of 1.5 min keeping constantly 5 %B, increasing in 0.5 min to 30 %B followed by gradually increasing to 50 %B in 10 min and using a flow rate of 0.3 mL/min on a Acquity BioResolve column (450 Å, 2.7 µm, 2.1 mm x 150 mm, Waters) maintained at 70 °C. The mobile phases were 0.1% (v/v) TFA in water (A) and 0.09% (v/v) TFA in acetonitrile (B). Detection of UV signals was performed at 214 nm. The mass spectrometer was equipped with an electrospray ionization source, which was used in positive ion mode with a scan range from m/z 500 to 4000, a collision energy of 6V and a cone voltage of 100 V.

### Statistical analyses

Statistical analyses were performed as indicated using GraphPad Prism version 7.03 (GraphPad Software, Inc., La Jolla, CA). Differences were considered to be significant when the probability value was <0.05.

## ACKNOWLEDGMENTS

We thank the Musculoskeletal Disease and the Diseases of Aging and Regenerative Medicine areas at Novartis Biomedical Research (BR) for their support, particularly Schwob Jürg, along with the rest of the BR community.

## AUTHOR CONTRIBUTIONS

K.HJ. and S.S2. (Summermatter Serge) designed *in-vitro* experiments and animal studies, respectively, analyzed results and interpreted data. H.F. performed sequence analyses and designed proteins. H.F. and S.S1. (Seddik Sabrina) planned and executed recombinant protein purification, analytics, and data interpretation. H.C. performed mass spec analyses. F.A., B.F., C.C.A., K.M., P.N.C., M.A., H.L., and A.M, conducted *in-vitro* experiments and/or contributed to their analyses. M.S., B.J. and P.E. performed animal studies. S.M. and L.X. run and analyzed biophysical assays. V.A., G.G. and C.P. prepared reagents for Retrogenix and readout the data. A.D., A.C., K.S., P.E. and D.B.C. performed FACS sorting of satellite cells and analyzed the transcriptomics data. L.C., D.A. and A.N conducted the histological experiments. R.J. and L.S. analyzed transcriptomic datasets from *in-vitro* experiments, animal studies and human trials. B.A1. (Bepperling Alexander) performed analytical ultracentrifugation experiments and their analyses. W.L. and B.A2 (Burget Aurelia) conducted cloning and expression, respectively. R.D. provided clinical application guidance throughout the project and was involved in conceptualization. K.M. provided resources and strategic oversight. S.S2. supervised the project, conceptualized the results and wrote the manuscript. All authors received and approved the final document.

