## Supplementary Material for "Exercise-Induced Myostimulin Enhances Muscle Function in Health and Disease"

### 1. Supplementary Methods

#### Sequence Analyses and Visualization

Sequence alignments, secondary structure prediction and the sequence logo were generated using Jalview (v. 2.10.5) and JPred4 [1,2]. Charge, hydropathy and intrinsic disorder propensity were calculated as previously described[3-5] and visualized using Wolfram Mathematica (v. 14.1). The BLAST search to obtain the consensus sequence of the repeat motif was performed on UniProt using the default settings. Sequences were filtered for ≥50 % identity, the presence of a signal peptide and a length of up to 200 residues. Isoforms from the same species were removed (**Supplementary Table 1**).

#### UV Fluorescence Spectra and Irradiation

A Horiba FluoroMax4 spectrofluorometer, operated by FluorEssence 3.9 software, was used to record fluorescence emission spectra, and to irradiate protein samples. Both types of experiments were carried out at 20 °C. To collect spectra, 60 µl sample were placed in a 3x3 mm Hellma Ultra-Micro QS quartz cell. Ten emission spectra were recorded for each sample, scanning from 330 nm to 500 nm (5 nm bandpass) in 17 s, with the excitation monochromator set to 310 nm (2 nm bandpass). For irradiation, 2.4 ml of protein solution were placed in a 1x1 cm Hellma QS quartz cell and irradiated for 30 min. During irradiation, the solution was continuously stirred at 750 rpm using a magnetic stir bar to minimize the inevitable inhomogeneity. The excitation monochromator was set to 275 nm, emission was monitored at 420 nm.

#### Analytical Size-Exclusion Chromatography (SEC) and Light Scattering

Analytical SEC was performed on a Waters Acquity UPLC system at 0.4 ml/min and 40 °C, using a Waters BEH200 (4.6x150 mm) column, with 50mM sodium phosphate, 400mM sodium perchlorate, pH 6.0 as running buffer. LC data were analyzed using Thermo Chromeleon software. For Multi-Angle Light Scattering (MALS) measurements, a Wyatt µDAWN and an Optilab UT-rEX were included in the SEC-UPLC flow path. MALS data were analyzed using Wyatt ASTRA software. Dynamic Light Scattering (DLS) experiments were carried out using an AvidNano w130i instrument and a 1.5x1.5 mm Hellma QS quartz cell. Measurements were carried out a 20 °C, and average correlation functions were calculated for 10x 10 s measurements.

#### Analytical Ultracentrifugation (AUC)

All AUC experiments were performed with an Optima analytical ultracentrifuge (Beckman, Krefeld, Germany) supplied with absorbance and interference optics. 480 µl of sample were loaded into assembled cells with sapphire windows and 12-mm path length charcoal-filled epon double sector centerpieces. The samples were analyzed at 50,000 rpm in an eight-hole Beckmann-Coulter AN50-Ti rotor pre-equilibrated at 20°C. Sedimentation was monitored at 230 nm, 260 nm, and 280 nm and continuously scanned with a radial resolution of 10 microns taking one replicate per time point. The total duration of the experiment was 14:30 hours resulting in 381 scans for each cell during that time. Data analysis was carried out using the software UltraScan 4[6]. Intensity data were subjected to two-dimensional spectrum analysis (2DSA) analysis with meniscus and iterative fitting followed by a parametrically constrained spectrum analysis (PCSA) with 100 Monte Carlo iterations. For 2DSA, the grid resolution for s was set to 64 (range 1–10 S), for f/f0 it was left at 64 (range 1–4). For PCSA, increasing sigmoid, decreasing sigmoid and straight-line parameterization were tested, returning "straight line" as best fit model. All UltraScan calculations were done in-house on a Ryzen128-1019 Gigabyte TRX40 AORUS PRO with an AMD Ryzen Threadripper 3990X 2.9 GHz 64 Core/128 Threads CPU. Samples were diluted to 0.1, 0.2 and 0.3 mg/ml, respectively, using PBS.

#### 2. Supplementary Results and Discussion

##### Recombinant Production of Myostimulin

Expression of myostimulin in *E. coli* led to the formation of inclusion bodies. As the protein was not expected to adopt a defined tertiary structure, purification was carried out under denaturing conditions followed by removal of the denaturant w/o addition of common refolding additives. When exploring different options to purify myostimulin, we observed visible precipitation of the purified protein below pH 5.5, corroborating the hypothesis that the actual pI of the protein is significantly higher than the calculated value of 3.6. We could purify WT myostimulin from *E. coli* inclusion bodies in sufficient yields and purity using reversed-phase (RP)-HPLC (which we carried out at neutral pH to avoid protein precipitation). As expected from the sequence analysis, purified WT myostimulin formed covalent dimers via an interchain disulfide bond. The fraction of covalent dimers was lower than 50 % immediately after purification and only slowly increasing over time even in absence of a reducing reagent.

Inclusion bodies of the myostimulin [C105W, F107L] variant, however, appeared to be less stable in the *E. coli* cytosol and was associated with a significant reduction in yields. Therefore, we generated a His-tagged NproEDDIE fusion of the [C105W, F107L] variant. The engineered swine fever virus protease NproEDDIE is an effective tool for forcing proteins, that are unstable or toxic to the host cell, into inclusion bodies[7]. During refolding, NproEDDIE is cleaving itself off and can be separated from the protein of interest by IMAC in flow-through mode. In case of myostimulin, we polished the IMAC flow-through fraction using RP-HPLC as we did for the untagged protein. The NproEDDIE fusion led to a significant improvement of myostimulin titers but also to additional challenges and losses during purification, as 100 % refolding and autocleavage of NproEDDIE could not be achieved. A possible explanation for the inefficient cleavage during NproEDDIE refolding is that rapid myostimulin oligomerization under non-denaturing conditions (see below) brought the fused NproEDDIE moieties in close proximity. The resulting high local concentration led to crosslinking of NproEDDIE folding intermediates or misfolded species via interchain disulfides.

##### Purified, recombinant myostimulin is intrinsically disordered and forms soluble oligomers under physiological conditions

Initially, we used basic sequence analysis to predict the protein's properties. The gene encoding C1ORF54 is unique and exclusively found in mammals. In the InterPro database, C1ORF54 orthologues are collectively termed Protein(/Domain) of Unknow Function, DFU4634. The amino acid sequence reported in the UniProt entry for human C1ORF54, Q8WWF1, which is derived from GenBank entry NM_024579.3, served as basis for the experiments described herein. Human C1ORF54 consists of 131 amino acid residues. The similarity-based annotation of the first 16 residues as signal peptide in UniProt was confirmed using SignalP (v. 4.1)[8]. We termed the mature C1ORF54 polypeptide, devoid of the signal peptide, myostimulin.

Myostimulin has a molecular weight of 13.2 kDa. It's high content in acidic residues results in a very low theoretical pI of 3.6 and an estimated net charge of -16 at pH 7.4 [5]. The protein's high negative charge in combination with its small size and low GRAVY score of -0.43 [3], raised the question if myostimulin might be at least in parts intrinsically disordered, instead of adopting a stable globular fold. In a first approach, we used the so called Uversky plot to estimate the likelihood for myostimulin to be "natively unfolded"[9]. For comparison, we also added yeast thioredoxin, a "native protein", and alpha-synuclein, a "natively unfolded" protein to the Uversky plot (**Supplementary Figure 1**). From its position in this plot, myostimulin could already be classified as "natively unfolded".

In a next step, we plotted charge, hydropathy (HP) and intrinsic disorder propensity (IDP) along the myostimulin polypeptide (Fig. 3a). This plot not only solidified the hypothesis that myostimulin may not adopt a defined tertiary structure but also revealed the protein's amphiphilicity: The N-terminus (residues 1—13) is highly negatively charged, the middle part (residues 14—100) is less charged but still hydrophilic, while the C-terminus (residues 101—115) is very hydrophobic. (The clustering of acidic residues at the N-terminus of myostimulin may shift the actual pI to higher values, as described for, e.g., alpha-synuclein[10].

The middle part of myostimulin can be further divided into three sections with distinct features: a tyrosine cluster (residues 14—27), a coiled-coil motif (residues 40—60, identified by Jalview/Jnet/LUPAS21, **Supplementary Figure 2**) and a region with high propensity for intrinsic disorder (residues 62—100, TOP-IDP score > 0.1, using a sliding window of 15 residues). Not unusual for intrinsically disordered regions, the latter is harboring a repeat motif (residues 73—82, identified by visual inspection upon sequence alignment with Jalview/MUSCLE, **Supplementary Figure 3** and **Supplementary Table 1**)) and 5 potential O-glycosylation sites (positions: 63, 66, 76, 81, 82; identified NetOGlyc-4.0[11]. The tyrosine cluster and the coiled-coil motif are likely involved in protein-protein interactions and therefore assumed to be essential for myostimulin function. (Myostimulin also contains a couple of potential Y, T and S phosphorylation sites, which would become relevant if the protein is translocating to the cytoplasm as part of its mechanism of action. The number of repeats found in myostimulin of different species varies between one and nine, with the vast majority, including human myostimulin, containing only two repeats. Noteworthy exceptions are mouse and rat myostimulin, containing four and six repeats, respectively. In about 2/3 of the species included in this analysis, the myostimulin repeats contain at least one methionine residue, while the two repeats of human myostimulin do not. This is interesting because we found human myostimulin to be sensitive to oxidation (See also discussion on Stability of Myostimulin below). As these methionine residues are not conserved, they are likely not essential for function but may act as internal radical scavengers, thereby modulating the molecules duration of action, suggesting opportunities for engineering.

The very hydrophobic and likely alpha-helical C-terminus (**Supplementary Figure 2**) could facilitate membrane but also protein-protein interactions. Considering the amphiphilicity of the myostimulin polypeptide nature in combination with its high propensity for intrinsic disorder, suggested that the purified protein may form micelle-like oligomers in solution by self-association via the hydrophobic C-terminus, as described for artificial, amphiphilic polypeptides before[12].

We thus used size exclusion chromatography combined with multi-angle light scattering (SEC-MALS) and dynamic light scattering (DLS), to assess the folding and potential oligomerization of recombinant myostimulin. From the SEC retention time and DLS data we estimated a hydrodynamic radius of 8 and 10 nm, respectively, for the myostimulin oligomers (**Supplementary Figure 4**). This radius would correspond to a 450–800 kDa globular protein/oligomer. However, the MALS data revealed a molecular weight of only about ~220 kDa, corresponding to 17 myostimulin chains per oligomer on average. The discrepancy between molecular weight and hydrodynamic radius of myostimulin in solution can be explained by a detergent-like oligomerization mode, driven by the protein's hydrophobic C-terminus, as predicted from the sequence analysis.

To further corroborate our hypothesis, we generated a myostimulin variant comprising only residues 1—100, termed ΔC. N-terminally His-tagged ΔC (MW = 11 kDa) was expressed in *E. coli* and could be purified in one step from the soluble fraction using IMAC. When analyzed by SEC-MALS, ΔC eluted at a retention time expected for a globular protein of 34 kDa, corresponding to a hydrodynamic radius of 2.8 nm; the data from the MALS detector, however, revealed a molecular weight of 11 kDa (data not shown). Hence, in contrast to WT myostimulin, ΔC is mainly monomeric. Due to the small size of ΔC and likely also because of its intrinsic chain dynamics, the quality of the DLS data we collected for this variant was low. Nevertheless, the hydrodynamic radius of about 2.5nm estimated from the DLS data is in good agreement with the 2.8 nm calculated from the SEC retention time, which matches the hydrodynamic radius expected for a natively unfolded coil of the same MW[13] Although we found ΔC to be 100% monomeric, we cannot rule out that parts of the peptide contribute to the oligomerization of full-length myostimulin.

To gain further insights into the oligomerization of myostimulin in solution, we used analytical ultracentrifugation. These experiments revealed a strong concentration-dependency of the derived hydrodynamic parameters like sedimentation coefficient, hydrodynamic radius, frictional ratio and molecular mass in the range of 0.1 to 3 mg/ml (**Supplementary Figure 5**). This hydrodynamic and thermodynamic non-ideality at quite low concentrations pointed already to an intrinsically disordered protein. This was confirmed by the fitted weight-average frictional ratio of 2.05 (**Supplementary Figure 6** C) at the lowest concentration (0.1 mg/ml) compared to a value of 1.2 for a well-folded globular protein or 1.5 for an asymmetric, but folded molecule like an antibody. The hydrodynamic radius (mean 7.6/median 8.1, (**Supplementary Figure 7**)) is in excellent agreement with our DLS data. Taken together, these two parameters explain the high apparent molecular mass (hydrodynamic radius) observed by SEC, as its calibration is performed with a set of globular protein standards. An intrinsically disordered protein migrates at a higher apparent molecular mass (retention time) than a well-folded one. The sedimentation coefficient (**Supplementary Figure 6** A and C) and molecular mass (**Supplementary Figure 6** B and D) distribution shows a series of four main peaks centered around 6 S and 200 kDa respectively. These four peaks correspond to a 9mer (~120 kDa), 12mer, 15mer and 18mer (~240 kDa) based on the monomer mass of 13.2 kDa (**Supplementary Table 2**). It is tempting to suggest a trimer as the basic building block of recombinant myostimulin, but it remains to be further elucidated if this is a biological relevant assembly mechanism or the entropically driven aggregation pathway of the recombinantly produced protein.

The first AlphaFold model of C1orf54 (AF-Q8WWF1-F1-model_v1, (**Supplementary Figure 8**)) became available in Juli 2021[14]. It confirmed a high degree of intrinsic disorder, the presence of an alpha-helix in the middle part (which matches the coiled-coil motif identified by Jalview/Jnet/Lupas21) and the alpha-helical conformation of the hydrophobic C-terminus. However, the AlphaFold model does not provide insights into the mechanism behind the oligomerization of myostimulin.

In summary, sequence analysis and experimental data suggest that myostimulin is an intrinsically disordered protein, that forms micelle-like oligomers in solution via its hydrophobic C-terminus. It remains to be shown if the observed oligomerization is an actual "feature" of myostimulin, or just an artifact of recombinant production and purification. Our highly active myostimulin variant [C105W, F107L] forms oligomers as well, although with lower aggregation numbers as observed for the WT (Rh = 6.6 nm/7.5 nm by SEC/DLS, MW = 150 kDa by MALS, corresponding to ~12 chains per oligomer on average; data not shown). We hope that the analytical data presented herein expedite further biophysical characterization of myostimulin and help to further elucidate its mechanism of action.

Since we produced myostimulin in *E. coli*, we cannot comment on the impact of potential post-translational modifications like the predicted O-glycosylation on the folding/disorder or oligomerization of protein. Such modifications may also impact the biological activity of myostimulin in one way or the other. However, the likely membrane association of myostimulin may render its production in eukaryotic expression systems challenging.

#### Stability of Myostimulin

As expected from the low pI and the clustering of acidic residues near the N-terminus of the protein, we observed the formation of insoluble aggregates of myostimulin, WT and variant at pH 5.5 and below. Freezing and thawing, even if done rapidly, not only led to aggregation but also a significant reduction in activity of the myostimulin fraction remaining in solution, while the protein remained active for several months if stored at 4 °C. We also noticed that the protein lost activity upon long exposure to light. Since modern fluorescent light sources, such as the ones used in a laboratory environment, may emit UV radiation, we performed a stress test using a fluorescence spectrophotometer. Indeed, we found that irradiation of myostimulin at 275 nm led to the formation of dityrosine (**Supplementary Figure 9**). It is noteworthy, that dityrosine formation has also been shown to occur in vivo, e.g. for antibodies in inflamed tissue and for alpha-synuclein[15,16]. Further investigations are required to clarify if dityrosine formation in vitro is just a reflection of myostimulin's general sensitivity to oxidation or if dityrosine formation also occurs in vivo. Since UV irradiation is just one way to trigger protein oxidation via free radical formation, we speculate that oxidation of myostimulin might be one of the mechanisms that limit the duration of the protein's effects in vivo. To preserve the biological activity of purified, recombinant myostimulin, 3 mM L-Met and 1 mM EDTA were included in the formulation, and the protein was stored at 4 °C, protected from light.

#### Rational design of the [C105W, F107L] variant

**Supplementary Figure 10** shows the alignment of the C-terminal region of 20 arbitrarily chosen orthologues of human myostimulin found in UniProt. In orthologues that lack the single cysteine residue present in human myostimulin a tryptophan residue is found in this position. In most of these cases, C-to-W is paired with a F-to-L mutation two residues downstream. We therefore engineered a human myostimulin variant harboring these two mutations, C105W and F107L.

#### 3. Supplementary Figures

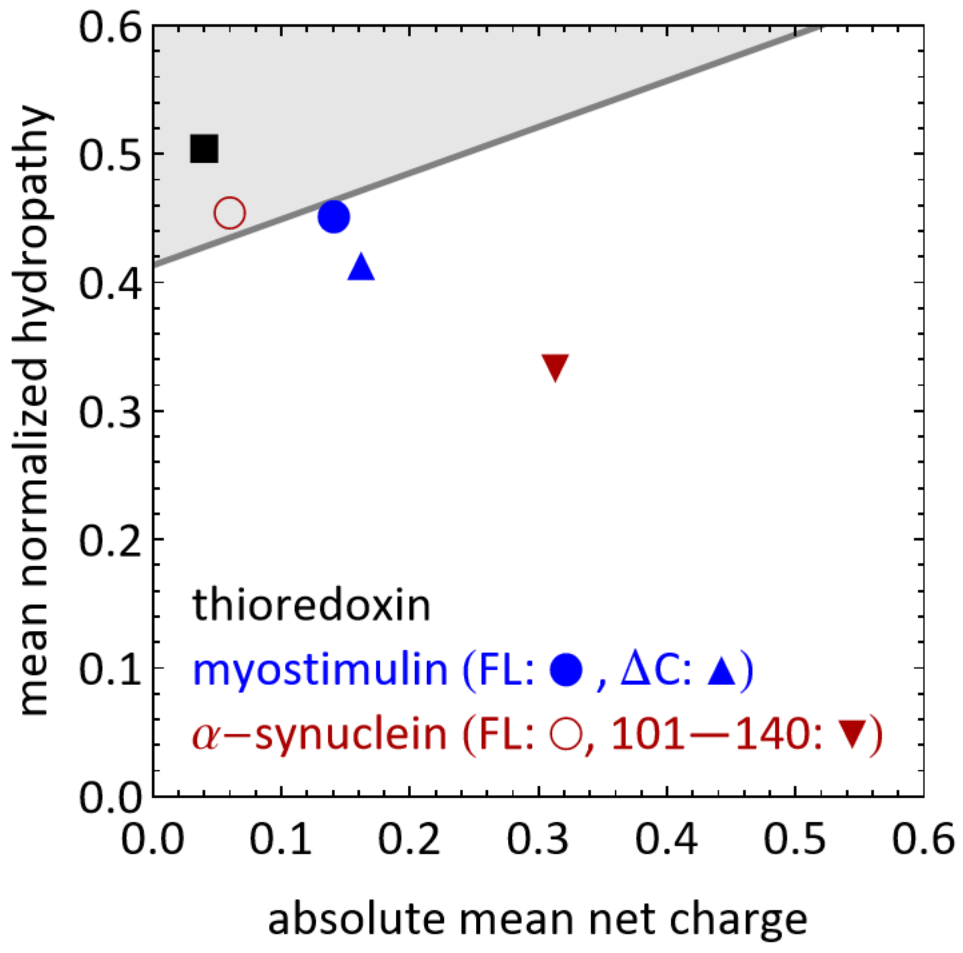


**Supplementary Figure 1:** Uversky Plot for the amino acid sequences of thioredoxin (black square), myostimulin (blue disc: full-length protein, blue triangle: residues 1—100) and α-synuclein (red circle: full-length protein, red triangle: residues 101—140).

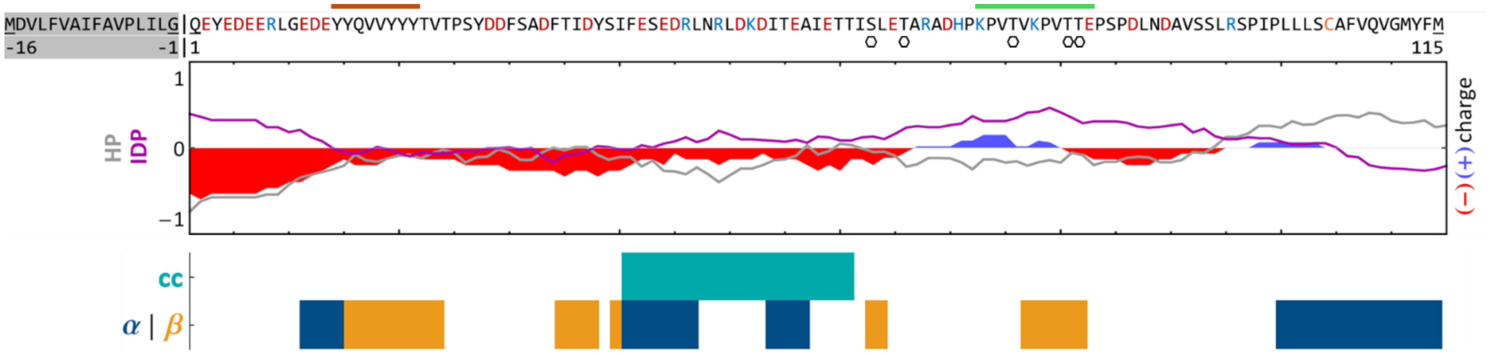

**Supplementary Figure 2:** Sequence analysis of human myostimulin. Top: Sequence of full-length C1ORF54 with its signal peptide marked in grey. Acidic residues are highlighted in red (Cys: orange), basic residues in blue. The orange bar marks the tyrosine cluster, the green bar the 2 repeats, and the black hexagons the potential O-glycosylation sites. Middle: Charge, normalized hydropathy (HP) and intrinsic disorder propensity (IDP) plotted using a moving window of 15 residues. Bottom: Secondary structure (α/β) and coiled-coil (cc) prediction by Jalview/Jnet.

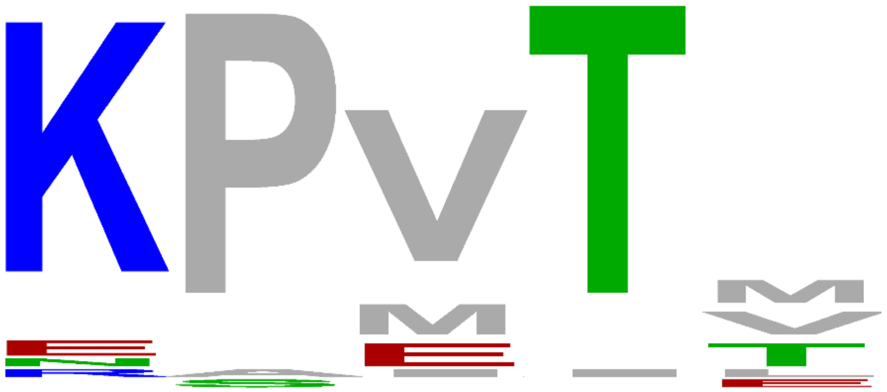

A)

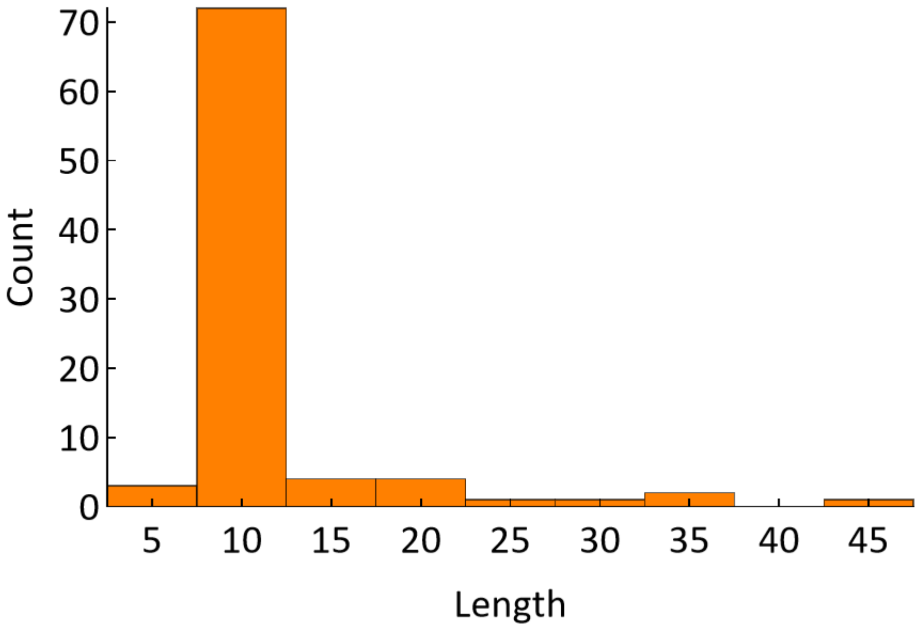


B)

**Q8WWF1|HUMAN/89-98**
KPVTV KPVTT
**Q148C7|BOVIN/88-97**
KPVIE KPVTM
**Q8R2K8|MOUSE/95-114**
NPVTT KPVTT EPVTT EPVTT

C)

**Supplementary Figure 3:** A) Sequence logo generated from the first two repeats of each myostimulin orthologue sequence (Supplementary Table 1). B) Length distribution of the repeat region C). Repeats for human, cattle and mouse.

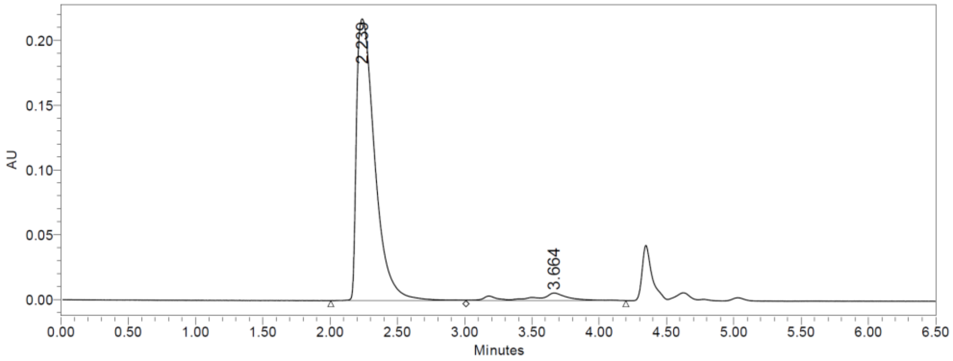


| Calibrant | RT/min |
|---|---|
| Thyroglobulin | 2.19 |
| IgG | 2.771 |
| holo-tranferrine | 3.079 |
| ovalbumin | 3.202 |
| carbonic anhydrase | 3.634 |
| aprotinin | 4.494 |

MW = 470 kDa
Calculated from calibration

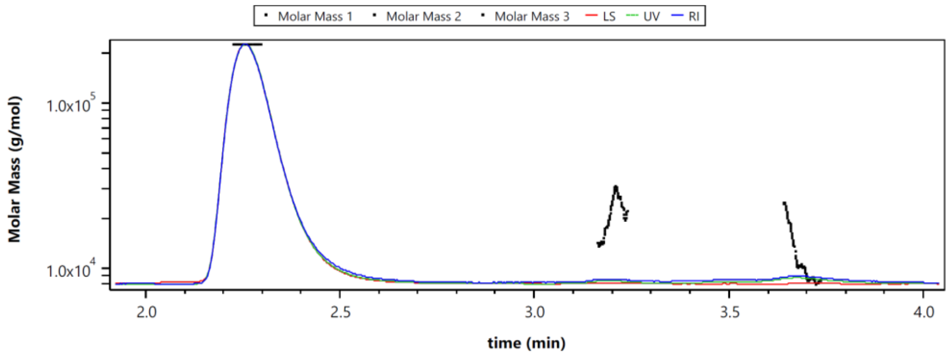


MW = 223 kDa
Calculated from MALS

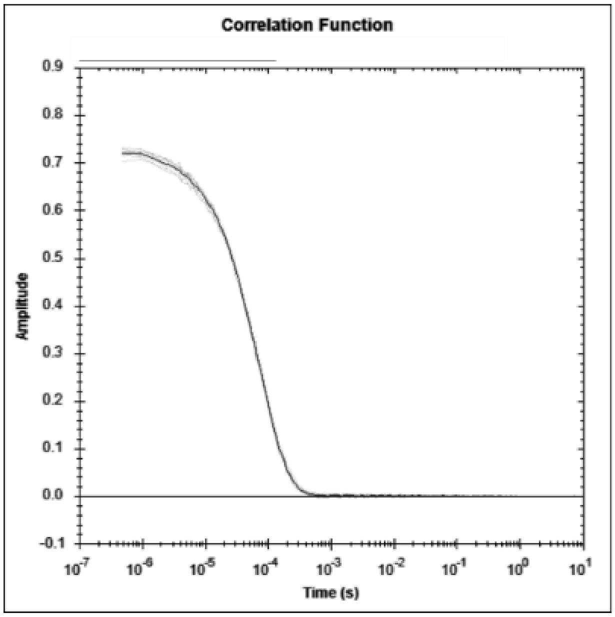


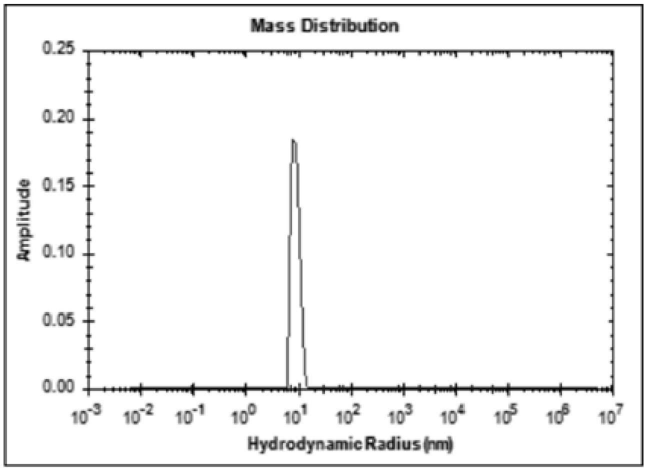


**Z - Av. Radius:** 9.86nm **Std. Deviation:** 5.61nm **MW model:** Globular Proteins

**Polydispersity:** 56.83 % **Pd. Index:** 0.323 **Remark:** -

| Peak # | Mean Rh (nm) | Mode Rh (nm) | Std. Dev. (nm) | Polydisp. (%) | Est. MW. (kDa) | Intensity (%) | Mass (%) | Volume (%) | Number (%) |
|---|---|---|---|---|---|---|---|---|---|
| 1 | 9.84 | 10.02 | 2.49 | 25.34 | 854.68 | 100.00 | 100.00 | 100.00 | 100.00 |

**Supplementary Figure 4:** Analysis of myostimulin by SEC-MALS (top two plots) and DLS (bottom).

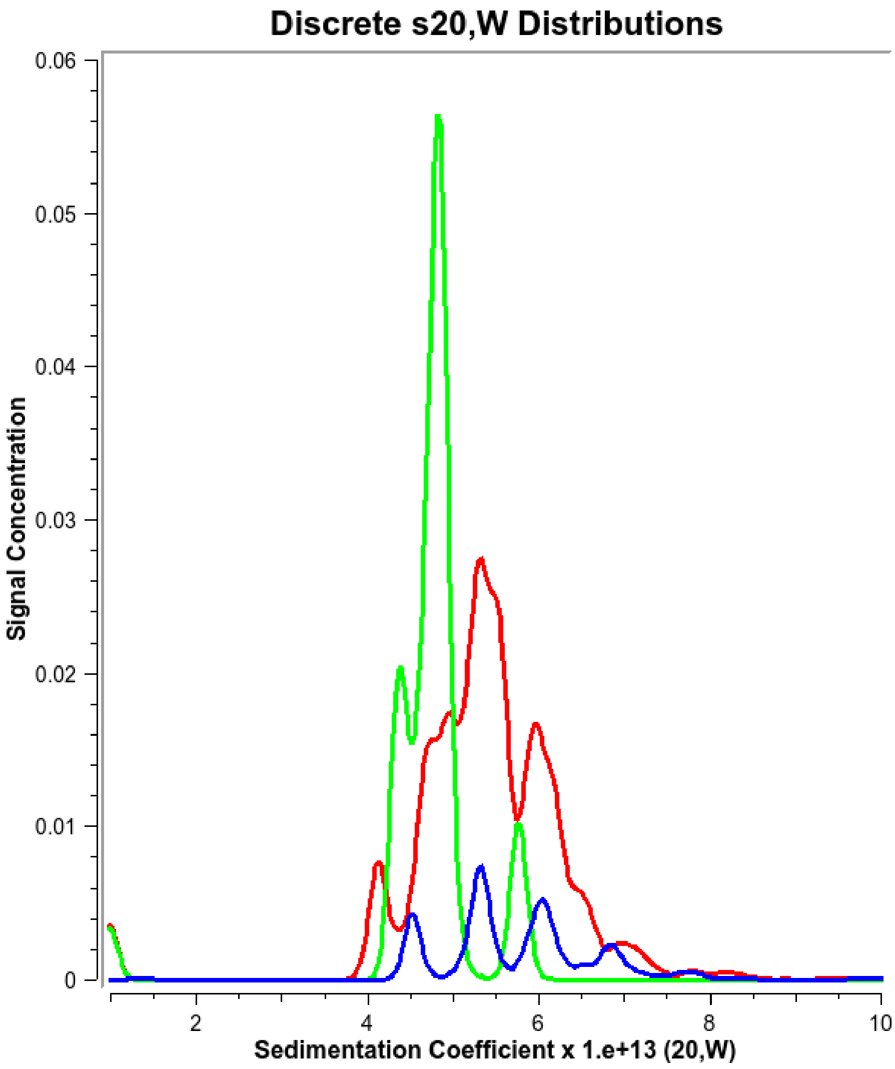


**Supplementary Figure 5:** Sedimentation coefficient distribution of 3 mg/ml (green), 1 mg/ml (red) and 0.1 mg/ml (blue) myostimulin in PBS. The decrease in the apparent weight-average sedimentation with increasing concentration (4.9 S (3 mg/ml), 5.5 S (1 mg/ml) and 5.8 S (0.1 mg/ml)) together with the artificial peak sharpening, particularly visible for the 3 mg/ml sample, indicate a strong hydrodynamic and thermodynamic non-ideality at already 1 mg/ml myostimulin concentration. Thus, the lowest concentration was chosen to determine the hydrodynamic properties of myostimulin in solution.

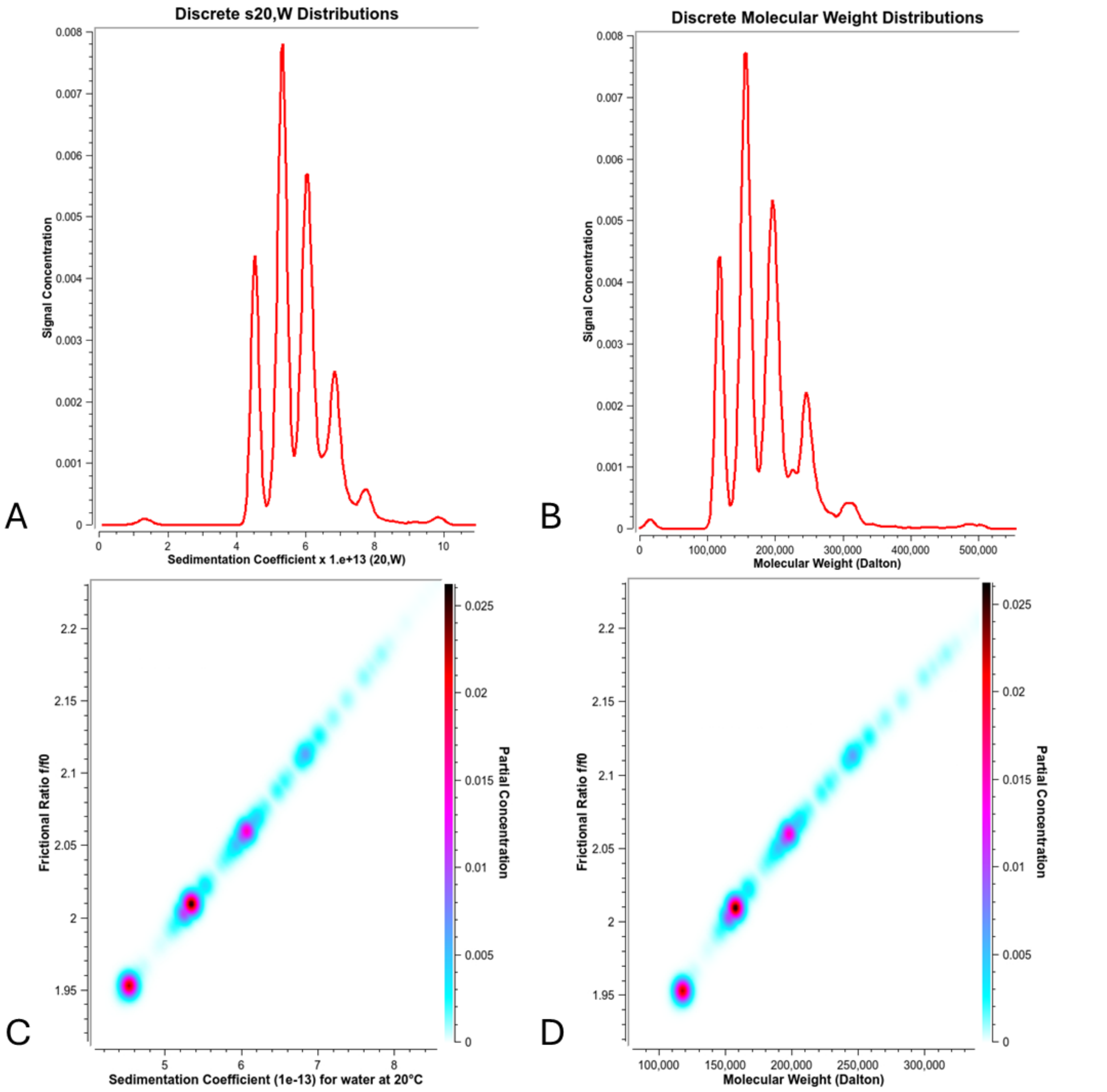


**Supplementary Figure 6:** Parametrically constrained spectrum analysis (PCSA-SL-MC) of 0.1 mg/ml myostimulin in PBS. Sedimentation coefficient (A) and molecular mass (B) distribution. Pseudo-3D representation of sedimentation coefficient vs. frictional ratio (C) and molecular mass vs. frictional ratio (D) indicating the randomly disordered secondary structure of myostimulin.

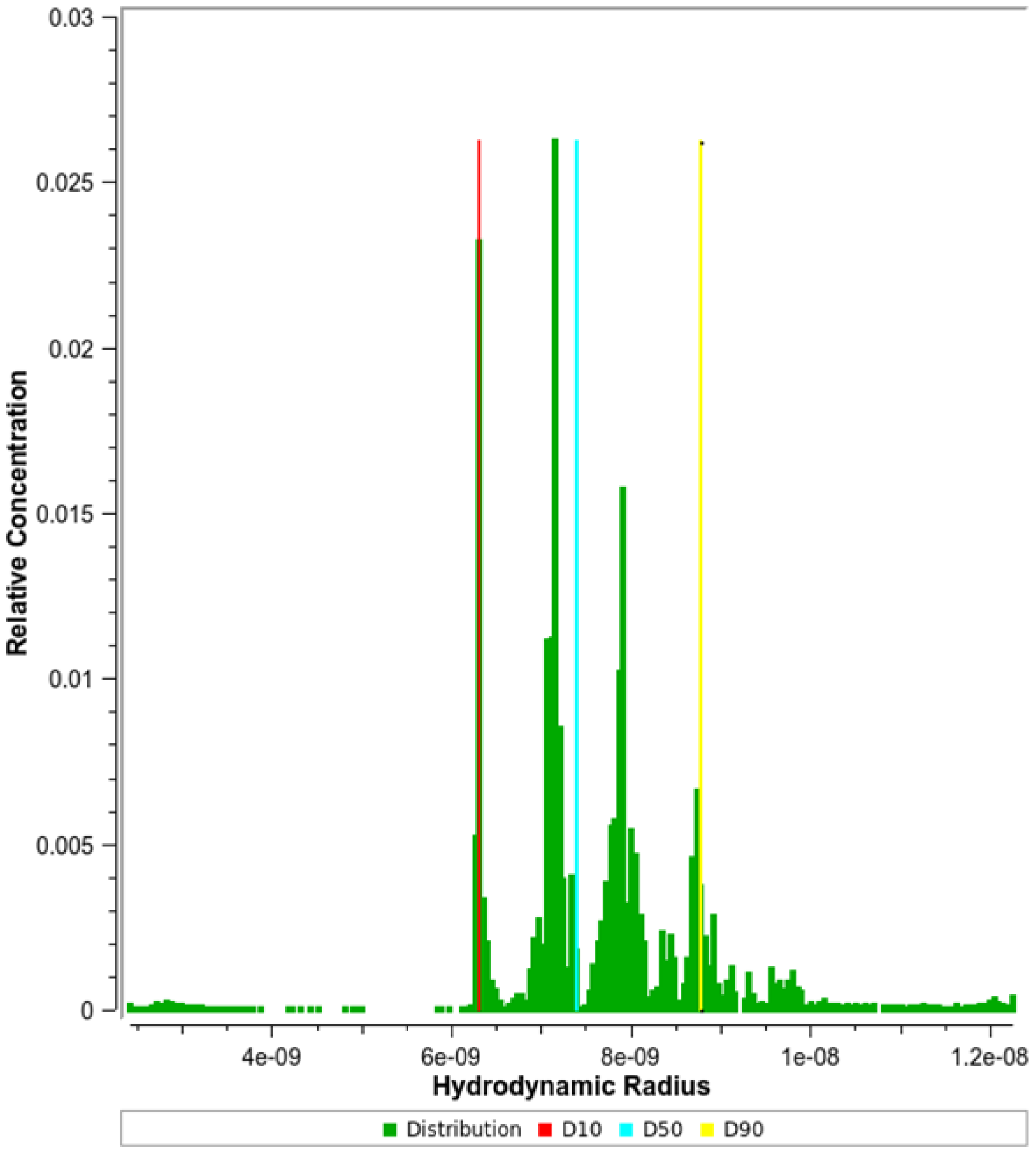


**Supplementary Figure 7:** Hydrodynamic radius distribution of 0.1 mg/ml myostimulin in PBS derived from the sedimentation coefficient and frictional ratio data of Supplementary Figure 6

**Supplementary Figure 6**.

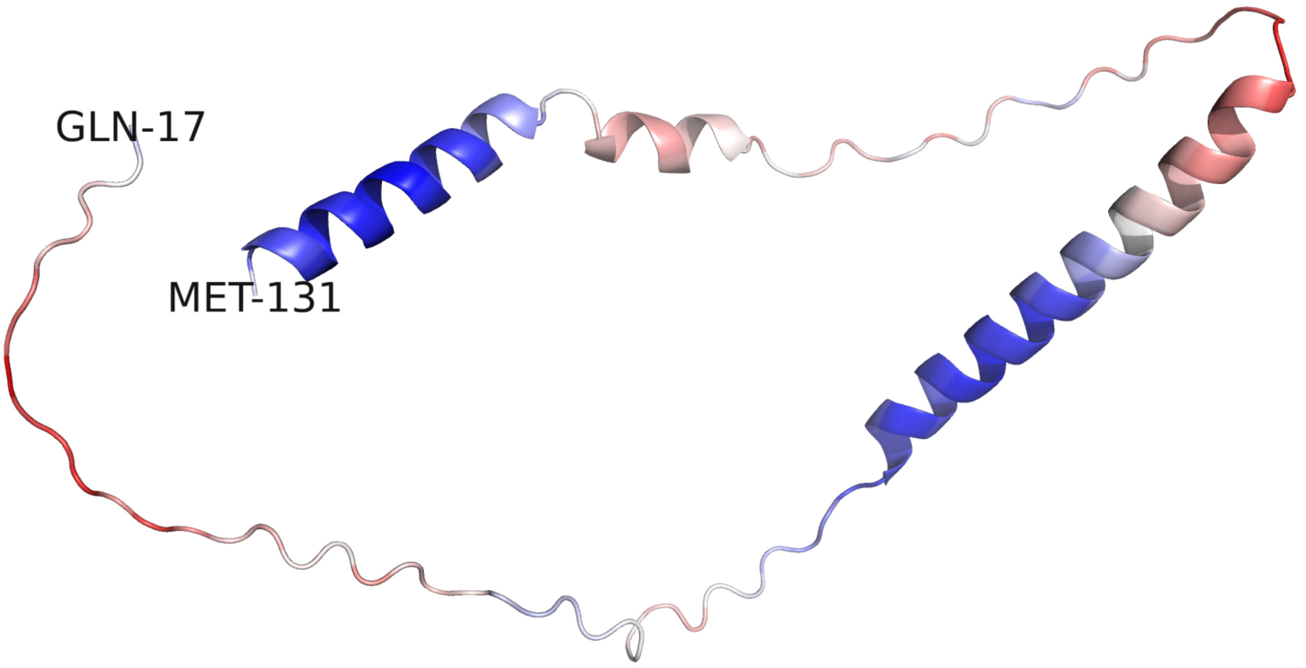


**Supplementary Figure 8:** AlphaFold1 model of myostimulin (C1ORF54, residues 17—131; blue/red—high/low confidence score).

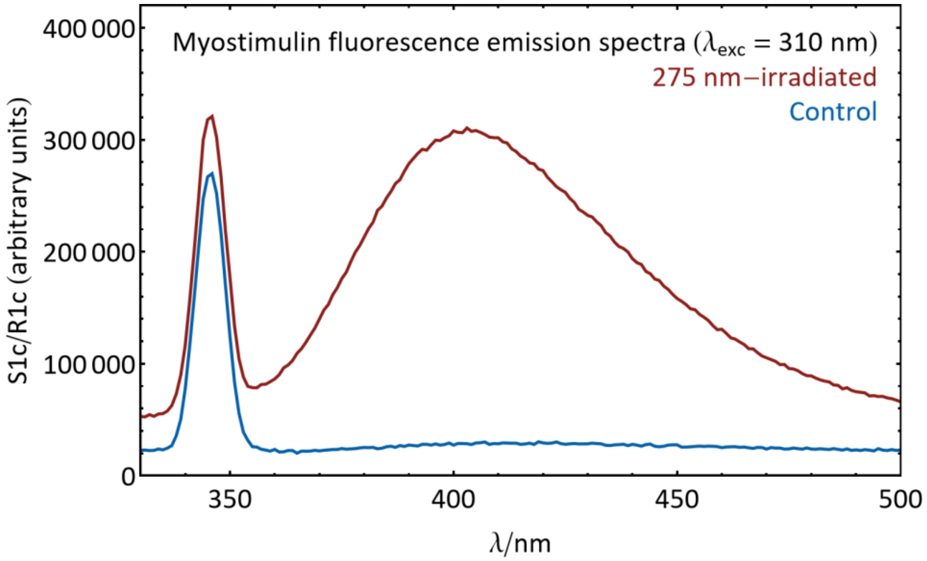


**Supplementary Figure 9:** Fluorescence emission spectra of myostimulin, illustrating the formation of dityrosine upon UV irradiation (right).

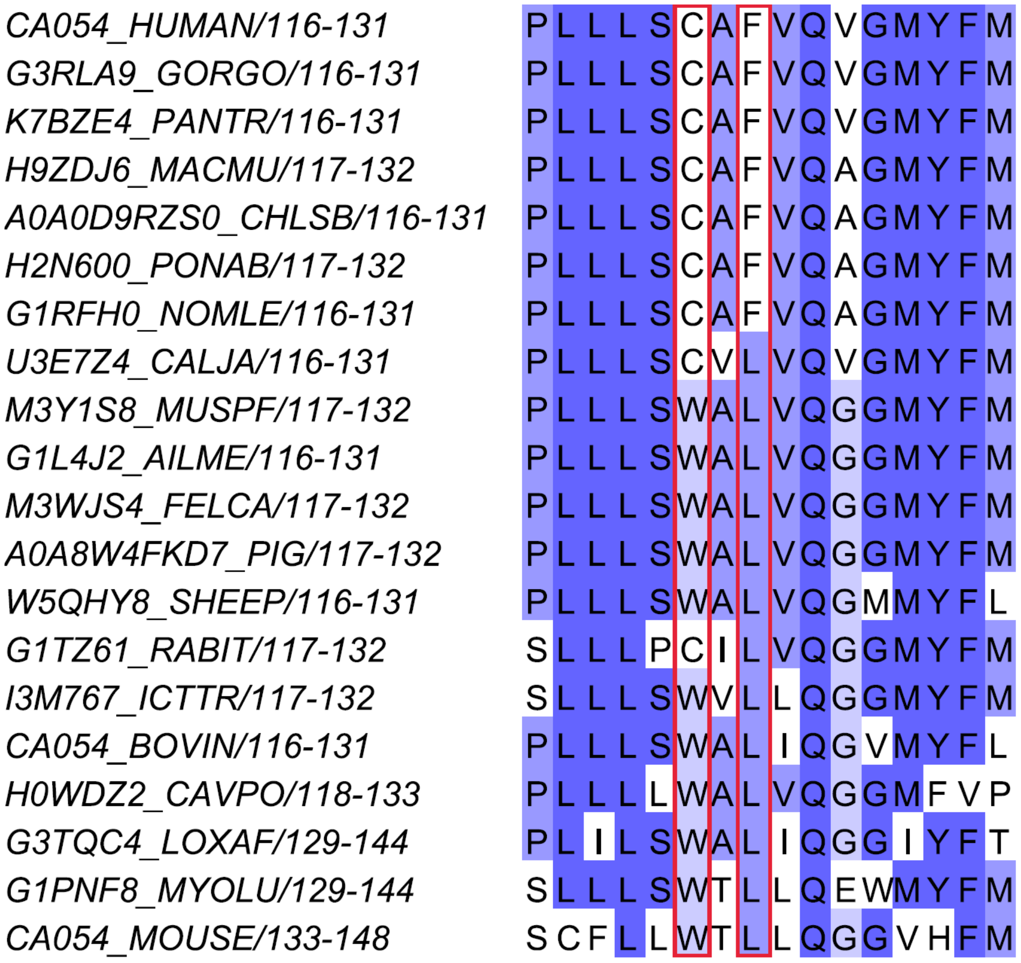


**Supplementary Figure 10:** Sequence alignment of the C-terminal region of 20 human myostimulin orthologues. The two positions selected for mutations, C105 and F107 of human myostimulin, are marked with red boxes. The numbering on the left is based on the full-length translation products and therefore includes the signal peptide.

### 4. Supplementary Tables

**Supplementary Table 1:** Sequences used to identify the repeat motif of myostimulin.

| Sequence ID and Segment Boundaries | Sequence of Repeat Region |
|---|---|
| sp\|Q148C7\|CA054_BOVIN/88-97 | KPVIE KPVTM |
| tr\|A0A6J0A5A5\|A0A6J0A5A5_ACIJB/90-99 | EPETL KPMTM |
| tr\|G3TQC4\|G3TQC4_LOXAF/89-103 | KPVTR KPVTM EPPSV |
| tr\|G1L4J2\|G1L4J2_AILME/89-98 | KPGTL KPMTV |
| tr\|A0A2K5CSA4\|A0A2K5CSA4_AOTNA/84-98 | KPVTV KPVIT KPVIT |
| tr\|A0A383ZSZ4\|A0A383ZSZ4_BALAS/89-98 | KPVTE KPVTM |
| tr\|A0A8B8W4J0\|A0A8B8W4J0_BALMU/89-98 | KPVTE KPVTV |
| tr\|U3E7Z4\|U3E7Z4_CALJA/89-98 | KPITV KPVIT |
| tr\|A0A3Q7N628\|A0A3Q7N628_CALUR/96-105 | KPETL KPMTM |
| tr\|A0A9W3EUK3\|A0A9W3EUK3_CAMBA/89-98 | KPVTV KHVTV |
| tr\|A0A8C0L845\|A0A8C0L845_CANLU/92-111 | KPETL KPMTVE PVTD SPLSV |
| tr\|A0A3Q0E5P3\|A0A3Q0E5P3_CARSF/89-98 | KPVTV KPVTT |
| tr\|A0A250Y9M9\|A0A250Y9M9_CASCN/90-94 | KPVTM |
| tr\|A0A8C3WER0\|A0A8C3WER0_9CETA/88-97 | KPVTM KPMTT |
| tr\|H0WDZ2\|H0WDZ2_CAVPO/89-98 | NPVTT KPMTM |
| tr\|A0A2K5RKN2\|A0A2K5RKN2_CEBIM/89-98 | KPVTV KPVTT |
| tr\|A0A2K5M432\|A0A2K5M432_CERAT/89-98 | KPVTL KPVTT |
| tr\|A0A8C2VP85\|A0A8C2VP85_CHILA/86-95 | KPVTT KPVTM |
| tr\|A0A0D9RZS0\|A0A0D9RZS0_CHLSB/89-98 | KPVTV KPVTT |
| tr\|A0A9B0T9H0\|A0A9B0T9H0_CHRAS/93-102 | KPVTM KPMTM |
| tr\|A0A2K5IUY7\|A0A2K5IUY7_COLAP/89-98 | KPVTV KPVTT |
| tr\|A0A2Y9MLT7\|A0A2Y9MLT7_DELLE/89-98 | KPVTE KPVTM |
| tr\|A0A1S3F271\|A0A1S3F271_DIPOR/102-111 | KPVTT KPITT |
| tr\|A0A2Y9LJN1\|A0A2Y9LJN1_ENHLU/89-98 | KPETV KPMTT |
| tr\|A0AA40LDU7\|A0AA40LDU7_EPTNI/89-133 | RPETV KPVTM EPVTM EPVTM EPVTM ESVTM GPVTM EPMTM EPVTM |
| tr\|A0A1S3WLV6\|A0A1S3WLV6_ERIEU/86-120 | EPVTV EPVTM NPLTT EPVIV EPVTM QPMTT KPVTK |
| tr\|M3WJS4\|M3WJS4_FELCA/90-99 | EPETL KPMTM |
| tr\|G3RLA9\|G3RLA9_GORGO/89-98 | KPVTV KPVTT |
| tr\|A0A9X9Q725\|A0A9X9Q725_GULGU/89-98 | KPETV KPMTT |
| tr\|A0A8B7PYY9\|A0A8B7PYY9_HIPAR/91-105 | KPVTV KPVKP VTMET |
| sp\|Q8WWF1\|CA054_HUMAN/89-98 | KPVTV KPVTT |
| tr\|I3M767\|I3M767_ICTTR/89-98 | KPVTM KAVTM |
| tr\|A0A2U3XAL8\|A0A2U3XAL8_LEPWE/89-93 | KPETM |
| tr\|A0A340XQF6\|A0A340XQF6_LIPVE/89-98 | KPVTE KPMTM |
| tr\|A0A667GVS5\|A0A667GVS5_LYNCA/90-99 | EPETL KPMTM |
| tr\|A0A485PRS5\|A0A485PRS5_LYNPA/90-99 | EPETL KPMTM |
| tr\|A0A2K5W5Q2\|A0A2K5W5Q2_MACFA/89-98 | KPVTV KPVTT |
| tr\|F7FDI6\|F7FDI6_MACMU/89-98 | KPVTV KPVTT |
| tr\|A0A2K6D1U8\|A0A2K6D1U8_MACNE/89-98 | KPVTV KPVTT |
| tr\|A0A2K5XET2\|A0A2K5XET2_MANLE/89-98 | KPVTV KPVTT |

| | |
|---|---|
| tr\|A0A5E4BSU0\|A0A5E4BSU0_MARMO/89-98 | KPVTV KAVTM |
| tr\|A0A3Q0DED2\|A0A3Q0DED2_MESAU/89-98 | NPVTT RPVST |
| tr\|A0A8B7GJ44\|A0A8B7GJ44_MICMU/89-98 | KPITV KPVIM |
| tr\|A0A7J8CPL5\|A0A7J8CPL5_MOLMO/89-98 | RPITV KPVLT |
| tr\|A0A8C6E9Q2\|A0A8C6E9Q2_MOSMO/89-98 | KPVTE KPMTM |
| sp\|Q8R2K8\|CA054_MOUSE/95-114 | NPVTT KPVTT EPVTT EPVTT |
| tr\|A0A8C6GNV9\|A0A8C6GNV9_MUSSI/95-104 | NPVTT KPVTT |
| tr\|M3Y1S8\|M3Y1S8_MUSPF/89-98 | KPETV KPMTR |
| tr\|G1PNF8\|G1PNF8_MYOLU/91-110 | RPVTV KSVTV EPVTM EPVTM |
| tr\|A0A7J7SPC6\|A0A7J7SPC6_MYOMY/91-110 | RPVTV KSVTV EPVTM EPVTM |
| tr\|A0A8C6QVV9\|A0A8C6QVV9_NANGA/89-98 | HPVTM KAVTM |
| tr\|A0A2Y9H381\|A0A2Y9H381_NEOSC/89-93 | KPETM |
| tr\|A0A341D6D7\|A0A341D6D7_NEOAA/89-98 | KPVTE KPVTM |
| tr\|A0A8C7BEL4\|A0A8C7BEL4_NEOVI/89-98 | KPETV KPMTR |
| tr\|G1RFH0\|G1RFH0_NOMLE/89-98 | KPVTV KPVTT |
| tr\|A0A2U3W7R3\|A0A2U3W7R3_ODORO/89-98 | KPETS KPMTM |
| tr\|A0A6J0Z0D7\|A0A6J0Z0D7_ODOVR/88-97 | KPVTE KPMTM |
| tr\|A0A8B7AMZ3\|A0A8B7AMZ3_ORYAF/90-99 | KPVTI KPVTM |
| tr\|G1TZ61\|G1TZ61_RABIT/90-99 | KPVTV KPTTM |
| tr\|A0AAD4URM3\|A0AAD4URM3_OVIAM/88-97 | KPVTE KPVTM |
| tr\|W5QHY8\|W5QHY8_SHEEP/88-97 | KPVTE KPVTM |
| tr\|A0A2R9C5F9\|A0A2R9C5F9_PANPA/89-98 | KPVTV KPVTT |
| tr\|A0A8C8Y118\|A0A8C8Y118_PANLE/90-99 | KPETL KPMTM |
| tr\|A0A8C9KQX8\|A0A8C9KQX8_PANTA/90-99 | KPETL KPMTM |
| tr\|K7BZE4\|K7BZE4_PANTR/89-98 | KPVTV KPVTT |
| tr\|A0A6J0DNB9\|A0A6J0DNB9_PERMB/91-115 | NPVTT RPARP ARPTR PTRPT RPATT |
| tr\|A0A8C9B598\|A0A8C9B598_PHOSS/89-98 | KPVTE KPVTM |
| tr\|A0A7E6CW67\|A0A7E6CW67_9CHIR/93-127 | KPVTM KPMTM GQMIT EPMTT ETTTT EPTTT EPVTM |
| tr\|A0A455B5N6\|A0A455B5N6_PHYMC/89-98 | KPVTE KPVKM |
| tr\|A0A7J7USH3\|A0A7J7USH3_PIPKU/90-99 | KPVTV KSVTV |
| tr\|H2N600\|H2N600_PONAB/89-98 | KPVTV KPVTT |
| tr\|A0A2K6EJU1\|A0A2K6EJU1_PROCO/89-98 | KPVTM KPVTM |
| tr\|A0A6P6GYB6\|A0A6P6GYB6_PUMCO/90-99 | EPETL KPMTM |
| tr\|A0A0G2JZ78\|A0A0G2JZ78_RAT/95-124 | NPVTT KPVTM KPVTT KPVTT KQVTT KQVTT |
| tr\|A0A2K6K864\|A0A2K6K864_RHIBE/89-98 | KPVTV KPVTT |
| tr\|A0A2K6PHS5\|A0A2K6PHS5_RHIRO/89-98 | KPVTV KPVTT |
| tr\|A0A7J8BC66\|A0A7J8BC66_ROUAE/89-103 | TSVTV KAVTM EPMEP |
| tr\|A0A6J3H0H1\|A0A6J3H0H1_SAPAP/100-109 | KPVTV KPVTT |
| tr\|A0A8D2CKB6\|A0A8D2CKB6_SCIVU/90-99 | KPITV KTVTM |
| tr\|A0A8C9QG07\|A0A8C9QG07_SPEDA/89-98 | KPVTV KAVTM |
| tr\|A0A673TMF7\|A0A673TMF7_SURSU/89-98 | KPETL KPMTV |
| tr\|A0A8W4FKD7\|A0A8W4FKD7_PIG/89-98 | KPVTM KPMTT |
| tr\|A0A8D2F7F6\|A0A8D2F7F6_THEGE/89-98 | KPVTV KPVTT |
| tr\|A0A2Y9E998\|A0A2Y9E998_TRIMA/100-109 | KPVTR KPATM |
| tr\|A0A2U4A3V3\|A0A2U4A3V3_TURTR/89-98 | KPVTE KPVTM |
| tr\|A0A8D2GKD9\|A0A8D2GKD9_UROPR/89-98 | KPVTV KAVIM |

| | |
|---|---|
| tr\|A0A452QZT0\|A0A452QZT0_URSAM/89-98 | KPGTL KPMTV |
| tr\|A0A384CGG9\|A0A384CGG9_URSMA/89-98 | KPGTL KPMTV |

**Supplementary Table 2:** Comparison of the fitted molecular mass of each of the four main peaks in Supplementary Figure 6 B and D against the theoretical molecular masses of one to 19 subunits of myostimulin.

| Subunits | calc. MW | MW (PCSA-MC) |
|---|---|---|
| 1 | 13221 | |
| 2 | 26442 | |
| 3 | 39663 | |
| 4 | 52884 | |
| 5 | 66105 | |
| 6 | 79326 | |
| 7 | 92547 | |
| 8 | 105768 | |
| 9 | 118989 | 118400 |
| 10 | 132210 | |
| 11 | 145431 | |
| 12 | 158652 | 159160 |
| 13 | 171873 | |
| 14 | 185094 | |
| 15 | 198315 | 198270 |
| 16 | 211536 | |
| 17 | 224757 | |
| 18 | 237978 | 245580 |
| 19 | 251199 | |

**Supplementary Table 3:** Overview of the hydrodynamic properties (weight-average) of myostimulin analyzed at 0.1 mg/ml in PBS.

| Parameter (Unit) | Value |
|---|---:|
| Sedimentation coefficient (S) | 5.80 |
| Frictional ratio | 2.05 |
| Molecular mass (kDa) | 210.00 |
| Hydrodynamic radius | 7.60 |

**Supplementary Table 4:** List of myostimulin binding proteins in Retrogenix, confirmatory protein array and transcriptomics data of satellite cells.

| Gene ID | UniProt ID | Classification | Myostimulin binding in Retrogenix | | Protein array | Receptor expression in satellite cells |
|---|---|---|---|---|---|---|
| | | | Replicate 1 | Replicate 2 | Average binding intensity (≥20 %) | (≥1000 counts) (Pax7 expression (reference)= 3529) |
| ICAM5 | Q9UMF0-1 | PM | | | | |
| PLXDC1 | Q8IUK5-1 | PM | | | | |
| **PLXDC2** | **Q6UX71-1** | **PM** | | | **21** | **1243** |
| NRXN3 | Q9HDB5-2 | PM | | | | |
| **IL6R** | **P08887-2** | **TS** | | | **30** | **2945** |
| CST6 | Q15828-1 | TS | | | | |
| GNRH2 | O43555-1 | TS | | | | |
| INSL4 | Q14641-1 | TS | | | | |
| INSL3 | P51460-1 | TS | | | | |
| SPINK6 | Q6UWN8-1 | TS | | | | |
| CPXM1 | Q96SM3-1 | TS | | | | |
| IGFBP3 | P17936-1 | TS | | | | |
| IFNW1 | P05000-1 | TS | | | | |
| IFNL1 | Q8IU54-1 | TS | | | | |
| LYZ | P61626-1 | TS | | | | |
| LYG2 | Q86SG7-1 | TS | | | | |
| KLK5 | Q9Y337-1 | TS | | | | |
| CLPS | P04118-1 | TS | | | | |
| DCN | P07585-1 | TS | | | | |
| ADA2 | Q9NZK5-1 | TS | | | | |
| NPPB | P16860-1 | TS | | | | |

| **Classification:** | **Binding intensity Retrogenix:** | Binding intensity Protein Array | Receptor expression in satellite cells |
|---|---|---|---|
| PM= Plasma Membrane | weak intensity | ≥20 % | ≥1000 counts |
| TS= Tethered secreted | medium intensity or above | | |
